# Expression of telomerase prevents ALT and maintains telomeric heterochromatin in juvenile brain tumors

**DOI:** 10.1101/718551

**Authors:** Aurora Irene Idilli, Emilio Cusanelli, Francesca Pagani, Emanuela Kerschbamer, Francesco Berardinelli, Manuel Bernabé, María Luisa Cayuela, Silvano Piazza, Pietro Luigi Poliani, Maria Caterina Mione

## Abstract

The activation of a telomere maintenance mechanism (TMM) is an essential step in cancer progression to escape replicative senescence and apoptosis. Paediatric brain tumors frequently exhibit Alternative Lengthening of Telomere (ALT) as active TMM, but the mechanisms involved in the induction of ALT in brain tumor cells are not clear.

Here, we report a model of juvenile zebrafish brain tumor that progressively develops ALT. We discovered that reduced expression of *tert* and increase in Terra expression precedes ALT development. Additionally, tumors show persistent telomeric DNA damage and loss of heterochromatin marks at chromosome ends. Surprisingly, expression of telomerase reverts ALT features. Comparative analysis of gene expression after the rescue of ALT with telomerase and analysis of telomerase positive paediatric brain cancers showed increase of telomeric heterochromatin and maintenance of telomere length compared to ALT tumors, with reduced expression of genes of the pre-replicative complex as hallmark. Thus our study identifies telomere maintenance mechanisms as major drivers of telomeric DNA replication and chromatin status in brain cancers.

## INTRODUCTION

Telomeres are nucleoprotein structures assembled at the end of eukaryotic chromosomes protecting them from fusions, degradation, and erroneous recombination events. In the absence of maintenance mechanisms, telomeres shorten at each cell division. Critically short telomeres trigger a DNA damage response, ultimately leading to an irreversible cell cycle arrest, known as senescence, or lead to apoptosis^1^. Telomerase is not expressed in somatic cells, which enter senescence upon a defined number of cell divisions^2^. In order to attain unlimited proliferative capacity, cancer cells must adopt telomere maintenance mechanisms. Most cancers reactivate the expression of telomerase a retrotranscriptase able to elongate telomeres through reverse transcription of its RNA subunit; however, a minority of cancers use alternative lengthening of telomeres (ALT), a mechanism based on homologous recombination between telomeres^3^. The molecular details of ALT activation remain to be defined^4^. ALT is found mostly in tumors with a mesenchymal origin (sarcoma) and in a subset of malignant paediatric brain tumors^5^, including High Grade Glioma (HGG, 51%), Diffuse Intrinsic Pontine Glioma (DIPG) (18%), Choroid Plexus Carcinoma (CPC) (22.6%), and Primitive Neuroectodermal Tumors (PNET) (11.6%)^6, 7^. HGGs represent approximately 3-5% of childhood brain tumors and include a heterogeneous group of rare but aggressive tumors^8^, which remains largely incurable, with the most aggressive forms being lethal within months. In contrast with adult glioma, mutations within the promoter region of the telomerase catalytic subunit, TERT, occur at a lower rate in paediatric tumors (3-11% compared to 55-83% in adult tumors)^9^. Indeed, paediatric gliomas show distinctive genetic mutations, which suggest that the mechanisms of tumor development and progression are different from those identified in adult brain tumors, where ALT develops in approximately 15% of cases, and associates with IDH1 mutations and better prognosis^10^. Exon sequencing of paediatric HGGs identified recurrent somatic mutations (K27M, G34R/V) in histone H3^11, 12^, and inactivating mutations in the Death-domain associated protein/Alpha thalassemia-mental retardation (DAXX/ATRX) genes, leading to DNA hypomethylation^13–15^. These findings suggest that telomeric chromatin plays an important role in ALT. Altered histone modifications in subtelomeric regions in association with ATRX loss result in deregulated telomere length and chromosomal instability, features that are often associated with ALT^4^. Notably, ALT activation can be inferred by a number of other features, including the presence of heterogeneous telomeres, with lengths ranging from very short to extremely long^16, 17^, the presence of circular extrachromosomal telomeric repeats (ECTRs) DNA, containing partially single-stranded telomeric CCCTAA repeats, also known as C-Circles, increased telomeric recombination, detected by the presence of telomere sister chromatid exchange (T-SCE) and formation of complexes of promyelocytic leukemia (PML) nuclear bodies with telomeres, known as ALT-associated PML bodies (APBs)^18^. Despite their heterochromatic nature, telomeres produce TERRA, a lncRNA playing an important role in telomere stability^19, 20^. TERRA expression has been shown to sustain activation of DNA damage responses at dysfunctional telomeres^21^ and TERRA transcripts are required for proper telomeric DNA replication in telomerase-positive human cancer cells^22^. Importantly, TERRA can form RNA:DNA hybrids or R-loops, at telomeres. Telomeric R-loops accumulate in ALT cancer cells, where they promote homologous recombination between chromosome ends, thereby sustaining ALT mechanisms^23^. Despite the established impact of telomere biology in cancer and the recent advancements on the role of chromatin structure and TERRA in TMMs in cancer, the molecular details that trigger ALT remain unclear. The study of ALT in cancer has mostly relied on ALT sarcoma cell lines^5^ and their telomerase positive counterparts, and only recently TMMs started to be evaluated in primary cancers^24^. Mouse models have been used extensively to study telomere biology^25^ and have enabled important advancements in the field. A caveat in the use of mice as a model for studying telomere biology is that mouse telomeres are considerably longer than human telomeres (50 kb vs 15 kb)^26^. In addition, telomerase continues to be expressed in most adult mouse tissues and organs, with the result that telomere shortening is not a problem for cellular lifespan in mouse. By contrast, zebrafish telomeres (15-20 kb) are relatively similar to human telomeres (10-15 kb) and, although telomerase is constitutively active in some organs, the expression of *tert* mRNA, the catalytic subunit of telomerase, telomerase activity and telomere length decrease drastically with age, similarly to human tissue; in addition, *tert* levels in the zebrafish brain are very low from early stages^27–29^.

Recently, we generated a zebrafish model of brain tumors based on somatic expression of human oncogenes^30^. In this model, brain tumors resemble the molecular mesenchymal subtype of glioblastoma^30^. Here we investigated the telomere maintenance mechanism adopted by brain tumor cells in this zebrafish model of brain cancer and found that brain cancer cells progressively develop ALT, upon reduction of telomerase expression and activity and increase of TERRA levels. Surprisingly, telomerase re-expression prevents the development of ALT and ameliorates cancer mortality and malignity. Comparison of gene expression between telomerase positive and ALT zebrafish brain tumors identifies reduced expression of genes of the pre-replicative complex and increased expression of genes driving heterochromatin deposition as hallmarks of the telomerase rescue. Analysis of telomerase levels and expression of 33 genes of the pathway “Activation of the pre-replicative complex” in 296 paediatric brain cancers from pedcBioportal (cbioportal.org) showed significant correlation; in ALT zebrafish and human juvenile brain tumors we found a significant increase of telomeric RPA+ foci, indicating stress responses to stalled replication forks, and an almost complete absence of H3K9_me3_ foci, indicative of a loss of heterochromatin.

## RESULTS

### Telomeres in zebrafish brain tumours are long and heterogeneous

We recently developed a zebrafish model of brain cancer, which is based on the injection of a plasmid encoding for UAS driven GFP-tagged HRAS^G12V^ in one-cell stage embryos of a transgenic line expressing the transactivator Gal4 in neural progenitor cells, *Et(zic4:Gal4TA4, UAS:mCherry)^hzm5^*, here called zic:Gal4. This leads to the integration and expression of the oncogene in a few neural progenitor cells, that become cancer-initiating cells^30^. In order to investigate telomere length in zebrafish brain tumors we used 1-2 month (m) old zebrafish with RAS-induced brain tumors, here called RAS, and performed terminal restriction fragment (TRF) analysis followed by Southern blot. Interestingly, RAS brain tumors showed highly heterogeneous telomere lengths as observed by the smeared bands detected on the TRF blot. In these samples, telomere size ranged from less than 6 kb to more than 23 kb in size (Figure 1a). By contrast, control brains showed a more compact band around 20 kb, indicating that more heterogeneous telomere lengths are present in zebrafish brain tumour cells compared to control brains (Figure 1a, S1a). Notably, telomere lengths of RAS brain tumors varied between samples and highly resemble the length of telomeres detected in U2OS cells (Figure S1a), which are known ALT positive human cancer cells.

**Figure 1.**
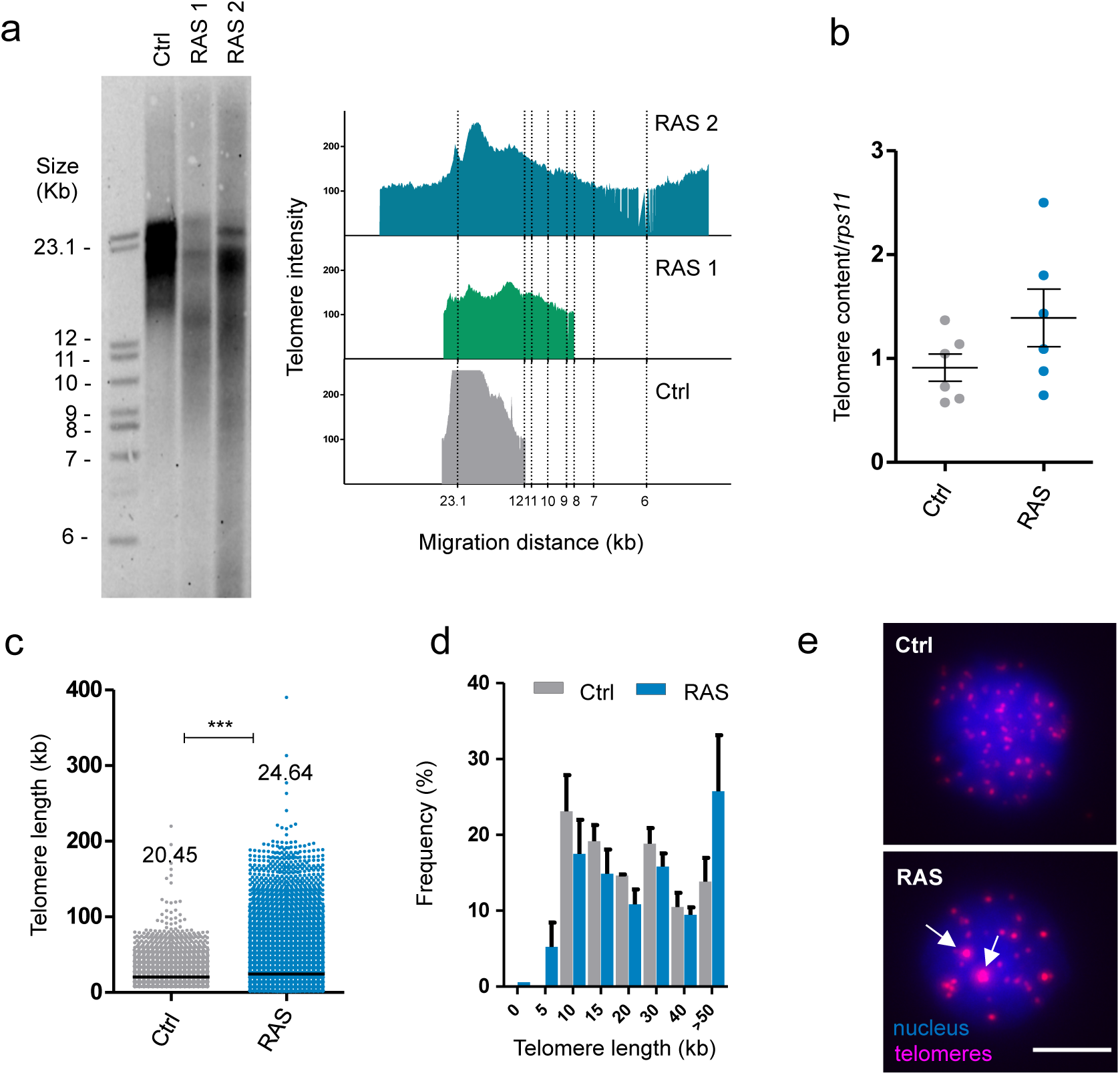
Zebrafish brain tumors have long and heterogeneous telomeres. **a)** Telomere length analysis via TRF in one control and two RAS tumors. The panel on the right shows TFR analysis obtained by graphing intensity of the signal versus DNA migration. **b)** Relative telomere content determined by telomere qPCR and normalized to the signal of a single copy gene (*rps11*) in controls (Ctrl, n = 7 brains) and RAS tumors (RAS, n=7 brains). Bars represent mean +/-s.e.m. **c)** Quantification of relative telomere length measured by Q-FISH and given a kb value based on the fluorescent intensity of L5178Y-S and L5178Y-R lymphocyte cells with known telomere lengths of 10.2 and 79.7 kb, respectively. See also figure S1b. Number of telomeres examined: Ctrl n= 3027, RAS = 9738. Data from three independent experiments were combined. Median values are reported on the graph. *** p <0.0001. Scatterplot bars represent median. **d)** Distribution of telomere length evaluated by Q-FISH in Control (grey n=2) and RAS tumors (light blue n=3). The very long telomeres (>30 kb) could represent telomeric clusters, an ALT feature. **e)** Representative fluorescent microscope images of Q-FISH analysis of control and RAS nuclei with ultrabright foci (white arrows). Scale bar: 5µm TRF: telomere restriction fragment; Ctrl: control; RAS: brain tumors

In order to confirm these results, we performed telomere qPCR analyses^31^ using genomic DNA extracted from RAS tumor samples or control brains. These experiments revealed an increase of telomere content in the RAS samples (Figure 1b), which is consistent with the results obtained from the TRF experiments. We then sought to evaluate differences in telomere length distribution at single-cell resolution, by performing Q-FISH experiments using fluorescently labelled telomere specific probe. In these analyses, we included as calibrators of single-telomere fluorescence intensity the L5178Y-S and L5178Y-R lymphocyte cell lines, which have telomere lengths of approximately 10 and 79 kb respectively^32^ (Figure S1b-c). Using this approach, Q-FISH experiments revealed a median telomeric length of 20.45 kb for control brains and 24.64 kb for tumour samples (Figure 1c). Interestingly, single-telomere fluorescence intensity distribution revealed the presence of very short telomeres (less than 5 kb) and very long telomeres (more than 50kb) only/preferentially in tumour cells as compared to control cells (Figure 1d). The presence of very long telomeres was also confirmed by the occurrence of ultra-bright telomeric foci^5^ (Figure 1e, lower panel).

Overall, these results indicate that brain tumor cells have highly heterogeneous telomeres as observed by TRF, telomere qPCR and Q-FISH analyses. Thus brain tumor cells may have activated a telomere maintenance mechanism, which is different from that of control brain cells.

### Telomerase is not involved in TMM in zebrafish brain tumours

Telomerase activity is tightly regulated during development and is re-activated in many tumours, where it is a critical determinant of cancer progression. To understand if the telomere length heterogeneity detected in zebrafish brain tumours is dependent on telomerase activity, we performed a quantitative telomeric repeat amplification protocol (Q-TRAP), an assay that allows to test the activity of the telomerase holoenzyme. By this analysis, we found a 2.5-fold reduction in telomerase activity in brain tumours (Figure 2a) compared to control brains of individuals of the same age. We next quantified the expression of the two components of the telomerase holoenzyme, the catalytic subunit, *tert,* and the template RNA, *terc,* through RT-qPCR (Figure 2b). *Tert* and *terc* transcript levels were significantly reduced in tumours (*tert* was reduced 1.7-fold; *terc* was reduced 1.9-fold) compared to control brains. In order to gain insight into the possible mechanism of *tert* downregulation, we analyzed the DNA methylation status of the zebrafish *tert* promoter in several brain tumours by performing chromatin immunoprecipitation (ChIP) of 5-methylcytosine enriched DNA sequences. Using EMBOSS CpGplot (https://www.ebi.ac.uk/Tools/seqstats/emboss_cpgplot/) we identified five regions in the *tert* promoter (GRCz11, chr: 19: 627,899-642,878; https://www.ensembl.org/Danio_rerio/), one of which has high CG content and is predicted to harbor a CpG island in position –1849 from the start of the gene coding sequence (Figure 2c, red arrows). We examined the methylation status of these five *tert* promoter regions in both control and brain tumours. We found that a general pattern of hypomethylation of the *tert* promoter was present in tumor samples as compared to control samples (Figure 2d). Each of the six tumours analyzed showed the same hypomethylated promoter status. The association between promoter hypomethylation and low expression has been reported also for human *TERT* in brain cancer^33^. This relation suggests that promoter hypomethylation is a general feature of negative regulation of *tert* expression, perhaps due to the location of the gene in a subtelomeric region (*CHR5:1253147-1295069*, GRCh38/hg38), both in human and in zebrafish (Figure 2c, inset). In summary, these results indicate that expression of *tert* is reduced in brain tumours and *tert* reduction associates with a hypomethylation status of the *tert* promoter that correlates with a significant decrease in the activity of telomerase. Overall, these findings suggest that telomeres in zebrafish brain tumours are not maintained by telomerase.

**Figure 2.**
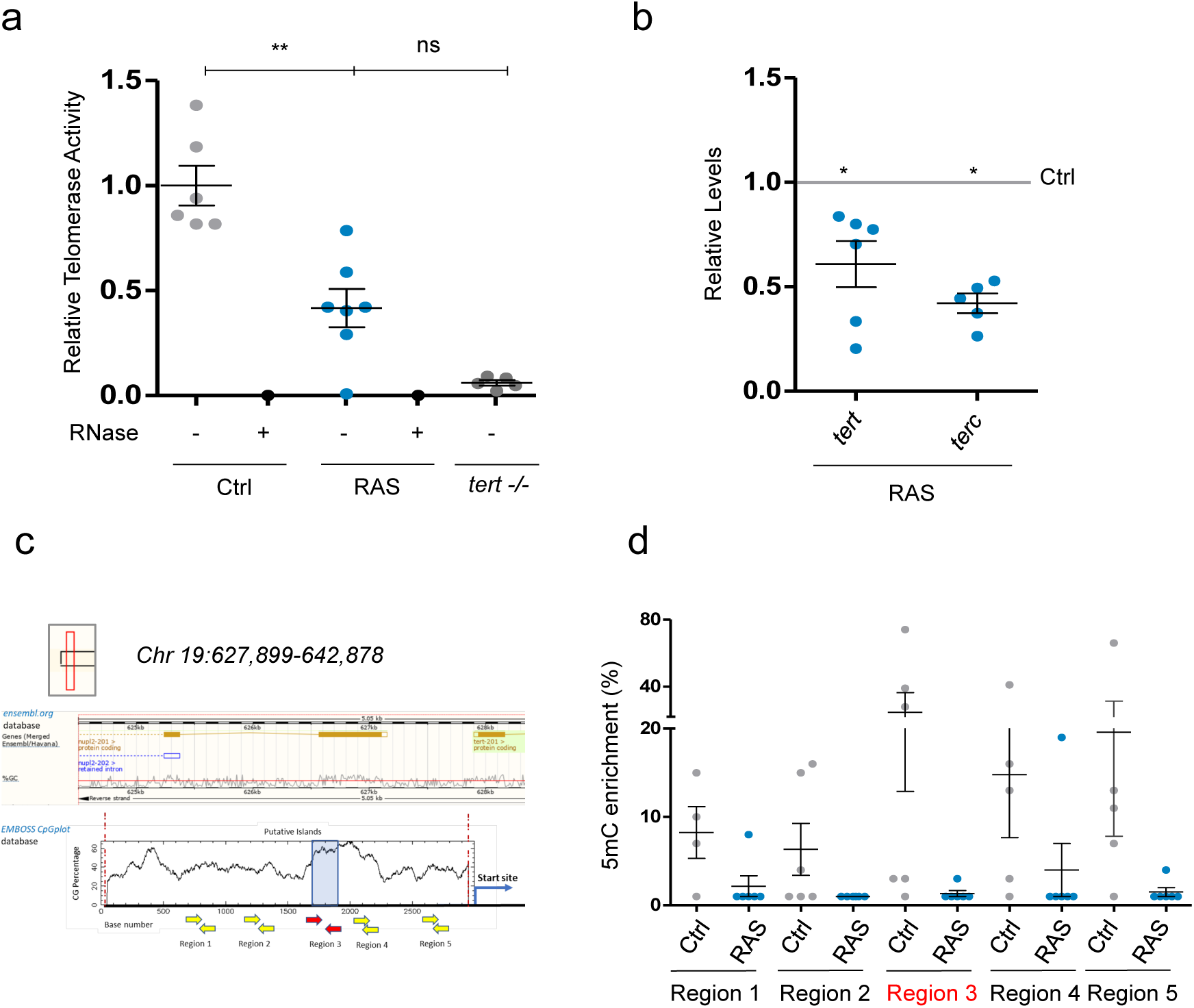
Telomerase is not involved in telomere maintenance in zebrafish brain tumours. **a)** Relative telomerase activity measured by Q-TRAP in control, RAS and *tert -/-* brains, using 1µg of protein extracts. RNase treatment (+) was used as a negative control to confirm the specificity of the assay, n =6;**p = 0.005. **b)** Expression of zebrafish *tert* and *terc* mRNA in brain tumors measured by RT-qPCR. Values were normalized to *rps11* mRNA levels and are relative to *tert* and *terc* expression in control brains (grey line set at 1.0) n =6; * p < 0.05. **c)** Schematic representation of the genomic region harbouring putative CpG island on the *tert* promoter, according to Ensemble (upper panel) and EMBOSS CpG plot (lower panel) databases. Position of primers used in ChIP experiments is shown as arrows. Red arrows show primers amplifying a putative CpG island. **d)** qPCR analysis of DNA CpG Methylation (5-Methylcytosine) status of the *tert* promoter in control and brain tumors of 2 month old fishes. Different regions of the promoter were analysed, red arrows indicate the position and primers for a putative CpG island. Values were normalized first to *rps11* and then to 5mC enrichment, with IgG enrichment set at 1.0. Bars in **a, b, e** represent mean +/-s.e.m. n =4 – 6. Ctrl: control brain, RAS: brain tumor

### Zebrafish brain tumours are ALT positive

In order to gain further insight into the telomere maintenance mechanism activated by the zebrafish brain tumor cells, we investigated whether ALT markers can be detected in tumors samples. We employed the C-Circle assay (CCA) to investigate the presence of C-Circles in zebrafish control and tumor samples, and, as additional controls, in two human cell lines, U2OS, an ALT cancer cell line, and HeLa, a telomerase-positive cell line^34, 35^. In these analyses, CCA products were detected by telomeric dot blot (Figure 3a) and telomeric qPCR (Figure 3b). Both methods showed that most of the tumours analyzed (10/13) were positive for C-Circles, with some variations in the levels of CCA products. Confirming the reliability of our approach, the amount of C-Circles detected in zebrafish tumor samples was similar to the amount of C-Circles detected in U2OS cells, while no positive signal was detected in HeLa or control brain samples (Figure 3a-b).

**Figure 3.**
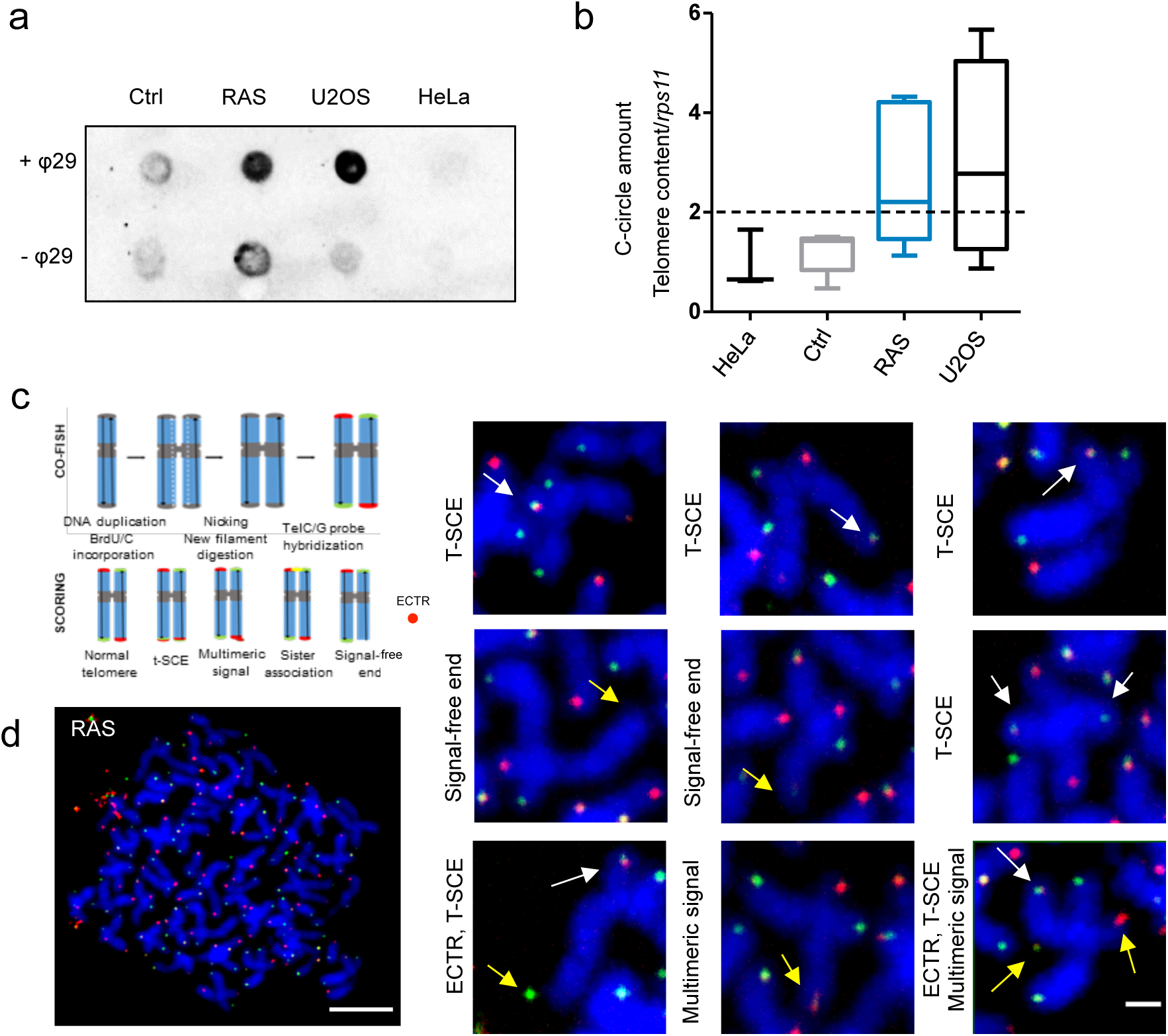
Zebrafish brain tumors are ALT. **a)** Representative C-Circle assay by dot blot in one control and one brain tumor compared with telomerase positive HeLa cells and ALT U2OS cells. Reactions without phy29 polymerase (-θ29) were included as a control. The assay was replicated 10 times. **b)** C-Circle assay quantified by telomere qPCR. Data are represented as amount of C-circles, normalized to telomere content (TC) and single copy gene (*rps11*). HeLa and U2OS were added as a reference. Ctrl n=5; RAS n=7. The dashed line indicates the level above which ALT activity is considered significant. Whiskers box plots represent median: min to max values. **c)** Schematic drawing to describe the procedure for 2-color CO-FISH and the interpretation of telomere status based on the signals. **d)** Two-color CO-FISH of a representative metaphase nucleus derived from a RAS brain tumor cell (scale bar 5µm). The right panels show details of telomeres with T-SCE (white arrows), signal free ends, multimeric signal and/or ECTR (yellow arrows) (scale bar 1µm). T-SCE: Telomere Sister Chromatid Exchange; ECTR: Extra-Chromosomal Telomeric Repeat; CO-FISH: Chromosome orientation FISH.

CO-FISH^36^ experiments (Figure 3c) revealed the occurrence of T-SCEs in brain tumor cells (Figure 3d, white arrows). In addition, the same analyses revealed the presence of telomeric defects, such as multimeric signals, signal free ends and ECTRs (Figure 3d, yellow arrows) on mitotic chromosomes of cells derived from zebrafish brain tumors.

We could not detect ALT-associated promyelocytic leukaemia (PML) bodies (APBs), due to the absence of a zebrafish ortholog of the PML gene^37^. ALT cancer cells express higher levels of the telomeric lncRNA TERRA than telomerase-positive cancer cells^13, 23, 38^. We investigated the expression of TERRA, which is known to localize to ALT-associated PML bodies^23^. TERRA is a RNA polymerase II transcript produced from subtelomeric regions towards chromosome ends^39^ (Figure 4a). To investigate TERRA expression, we performed RNA-FISH using TERRA-specific locked nucleic acid (LNA) probes^40^ in control and tumour cells (Figure 4b). By this analysis, we observed a significant increase in TERRA foci number and intensity in tumour cell nuclei versus controls (Figure 4c, d). The signal was abrogated by RNase treatment, confirming the specificity of the assay (Figure 4b, +RNase). In order to confirm these results, we performed RNA dot-blot analyses on total RNA extracted from RAS or control samples. These experiments confirmed an increase in TERRA levels in tumors as compared to control brains (Figure 4e), which is consistent with the RNA-FISH results. Finally, TERRA expression levels were quantified by performing quantitative reverse transcription PCR using total RNA obtained from control and brain tumours^33^ and compared to TERRA levels in HeLa and U2OS cells. By these analyses, we found that TERRA levels in tumours were almost 3-fold higher than in control brains (Figure 4f, left panel). As expected, TERRA levels were found higher in U2OS cells as compared to the telomerase-positive HeLa cells (Figure 4f, right panel). Overall, these findings thus indicate that our zebrafish model develops brain tumors, which are bona fide ALT.

**Figure 4.**
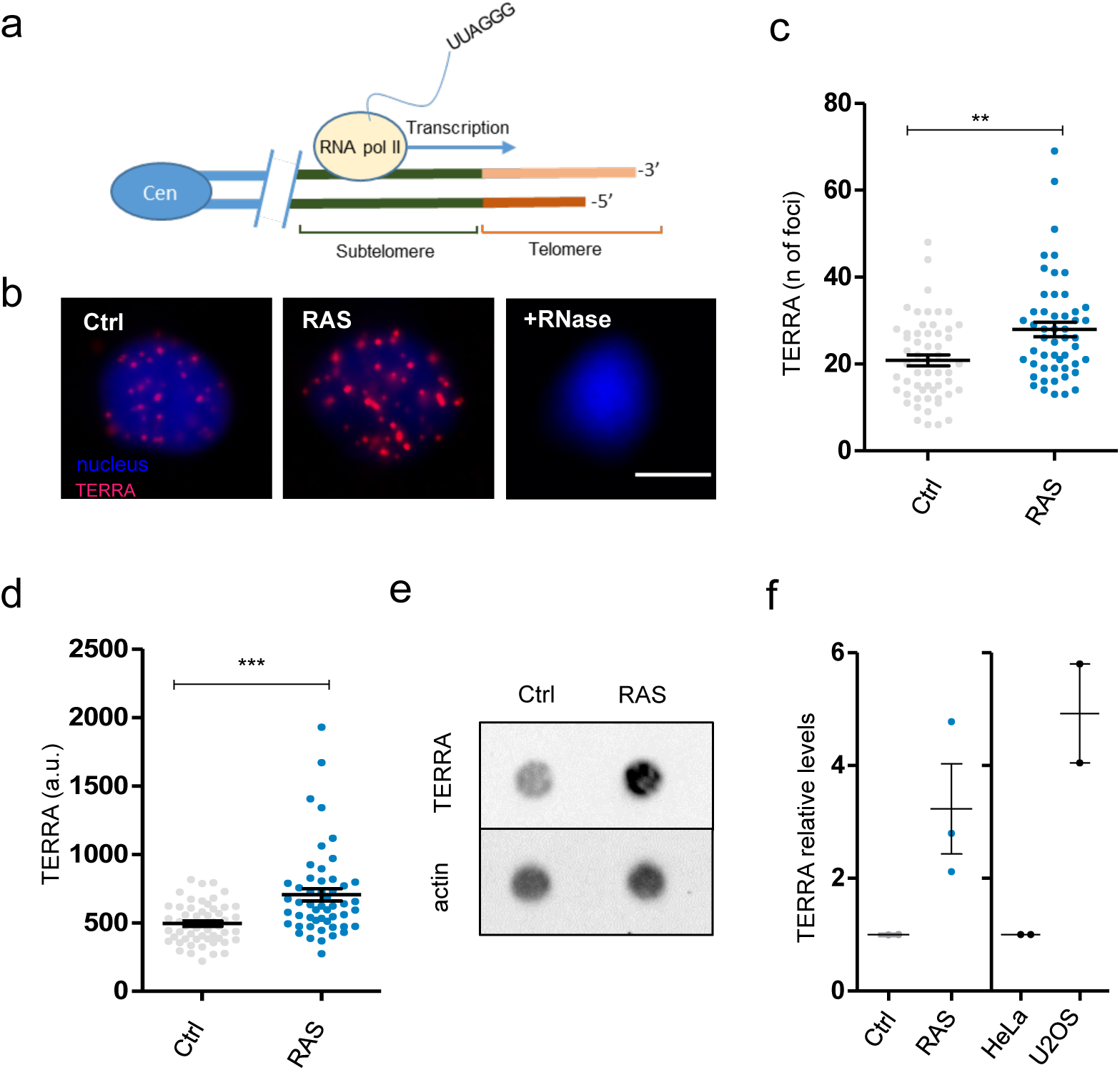
Zebrafish brain tumors are ALT and exhibit elevated TERRA expression. **a)** Schematic drawing describing the generation of TERRA from subtelomeric regions. Cen=centromere, RNA pol II =RNA polymerase II **b)** Representative pictures of TERRA RNA-FISH in cell nuclei (blue) from control (ctrl), tumor (RAS) and RNase treated tumor cells (+RNAse). TERRA foci are shown in magenta. Scale bar: 5µm **c)** Scatter plot of TERRA RNA-FISH expression measured as the number of foci per nucleus in Control (Ctrl, n =55) and tumor (RAS, n=51) nuclei. Representative data from one of three experiments are shown (**p =0.013). Bars represent mean +/-s.e.m. **d)** Scatter plot of TERRA RNA-FISH signals measured as fluorescent intensity per spot per nucleus in control (Ctrl) and RAS brains. Representative data from one of three experiments are shown (***p < 0.001). Bars represent mean +/-s.e.m. **e)** TERRA expression measured by dot blot from total RNA (500ng) of a control and a RAS brain tumor (upper panel); to control for RNA loading, hybridization with an actin RNA probe was performed (lower panel). **f)** qPCR analysis of TERRA expression in brain tumors and controls. Values were normalized first to *rps11* mRNA levels and then related to TERRA expression in control brains (n =3). TERRA expression levels were evaluated also in HeLa and U2OS cells (n=2 samples each) for comparison. Bars represent mean +/-s.e.m.

As in this zebrafish model, brain tumors are induced by the expression of the human oncogene RAS, we asked whether RAS expression may induce ALT in zebrafish. To answer this question, we used a different zebrafish model of cancer in which tissue-specific expression of RAS induces melanoma^41^ (Figure S2a). Detailed analyses of ALT hallmarks revealed that ALT did not develop in this model, in which the levels of *tert* expression were detected as 5-fold higher than in control skin (Figure S2b-c). These results indicate that HRAS^V^^12^ expression is not sufficient to activate ALT in zebrafish.

Similarly, we tested if ALT could be the preferred TMM of zebrafish brain cells when they form tumours. We generated cerebellar tumours using myristylated AKT (AKT) under the control of the zic:Gal4 promoter^30^ (Figure S2d) and we observed that neither ALT activity nor *tert* gene expression was changed compared to control cerebellar tissue (Figure S2e-f). Cerebellar tumours induced by AKT overexpression arise sporadically, mainly after four months (data not shown), when the fish are adults, as opposed to the RAS induced tumours that start to develop after 2-3 weeks^30^. Thus, the brain tumour model generated through the overexpression of oncogenic RAS in neural progenitors is the first in *vivo* GEM model of ALT brain tumours, and this feature is not due to the RAS oncogene and may be related to the juvenile age of the fish in the RAS model.

### A reduction of *tert* expression precedes the development of ALT

The model of brain tumour presented here allows the study of progression from a single cancer-initiating cell to a full tumour^30^. We investigated when ALT was activated during the progression of brain cancer, by performing CCA from the early stages (3dpf) to full tumour development (1m) (Figure 5a). This time interval corresponded to the entire larval period during which tumours grow progressively. In these experiments, we observed the presence of C-Circles above control levels starting from 20 dpf (Figure 5a). During the same period, we also studied the changes in *tert* expression (Figure 5b) and TERRA levels (Figure 5c-d). Lower levels of *tert* expression compared to control brains of the same age were observed throughout tumor development and a further decrease of *tert* levels preceded the increase of C-Circles at 20 dpf (Figure 5b). In particular*, tert* expression levels decrease in brains developing tumours to reach 1.7-fold less than controls at 1m (Figure 5b), when the tumours were fully formed.

**Figure 5.**
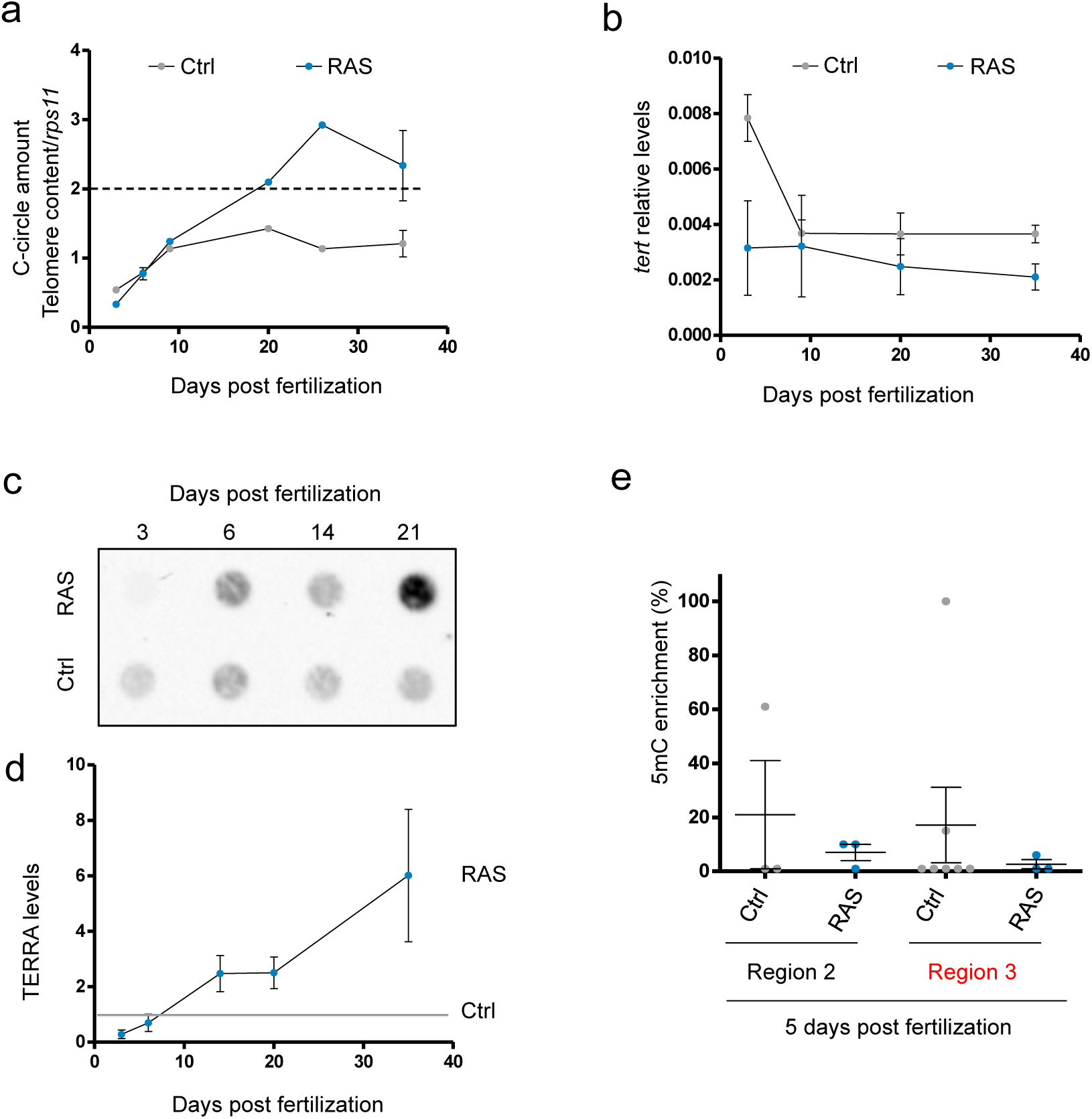
Development of ALT is preceded by a reduction of *tert* expression. **a)** C-Circle assay measured by telomeric qPCR during tumor development in control and RAS brains. Data are represented as CCA amount normalized to telomere content (TC) and single copy gene (*rps11*). The dashed line indicates the level above which ALT activity is considered significant Bars represent mean +/-s.e.m. **b)** RT-qPCR analysis of *tert* expression during tumor development. The data were normalized first to *rps11* mRNA levels and are expressed as 2 ^(-ΔCt)^. Bars represent mean +/-s.e.m. The experiment was replicated almost three times for each time points. **c)** Representative dot blot of TERRA levels during tumor development (500ng of total RNA was spotted for all samples) and **d)** quantification of three independent experiments. Background was removed and values were normalized to the levels of TERRA in controls of the same larval stages (grey line). **e)** qPCR analysis of DNA methylation (5-Methylcytosine, 5mC-ChIP) status of the *tert* promoter in 5dpf control (n = 3-5) and RAS (n = 3) fish larvae. Two regions of the promoter (see figure 2c) were analysed, the red color of region 3 indicates a putative CpG island. Values were normalized first to *rps11* and then to 5mC vs IgG enrichment, which is set at 1.0.

Furthermore, TERRA levels positively correlate with ALT activity and an increase of TERRA expression compared to control was already detected at 14dpf (Figure 5c-d), just before the increase in C-Circle levels.

We then examined the methylation status of the *tert* promoter at 5 dpf. We considered two of the most representative regions upstream of the transcriptional start site of the *tert* gene (Figure 2c, regions 2 and 3). We found hypomethylation of the *tert* promoter in RAS compared to controls already at 5dpf (Figure 5e), suggesting that the expression of *tert* in the brain was negatively modulated from the first days of RAS expression. Moreover, we found abnormalities in telomere signals in metaphases already at 5dpf (Figure S3a-b, yellow arrows).

In summary, these results suggest that a reduction of *tert* expression and an up-regulation of TERRA levels precede tumour formation and the activation of ALT.

### Overexpression of functional telomerase prevents ALT development

To evaluate if ALT activation could be prevented by maintaining high levels of telomerase, we increased *tert and terc* levels with transgenesis. To this aim, we first generated two stable transgenic lines, *tg(10xUAS:tert)* and *tg(10xUAS:terc)* (Figure S4a-b). The identification of F1 embryos expressing the transgenes was possible thanks to two different transgenesis markers which allowed us to combine the two transgenic lines and follow the inheritance of the markers under the fluorescent stereomicroscope (Figure S4c). Then we generated a triple transgenic line were *tert* and *terc* were overexpressed in neural progenitor cells (zic:Gal4+ cells) and we induced the development of brain tumours by injection the plasmid *UAS:GFP-HRAS^V12^*at one-cell stage. The generated brain tumors, indicated as RAS-Tert (Figure 6a), were compared with RAS (only) tumors.

**Figure 6.**
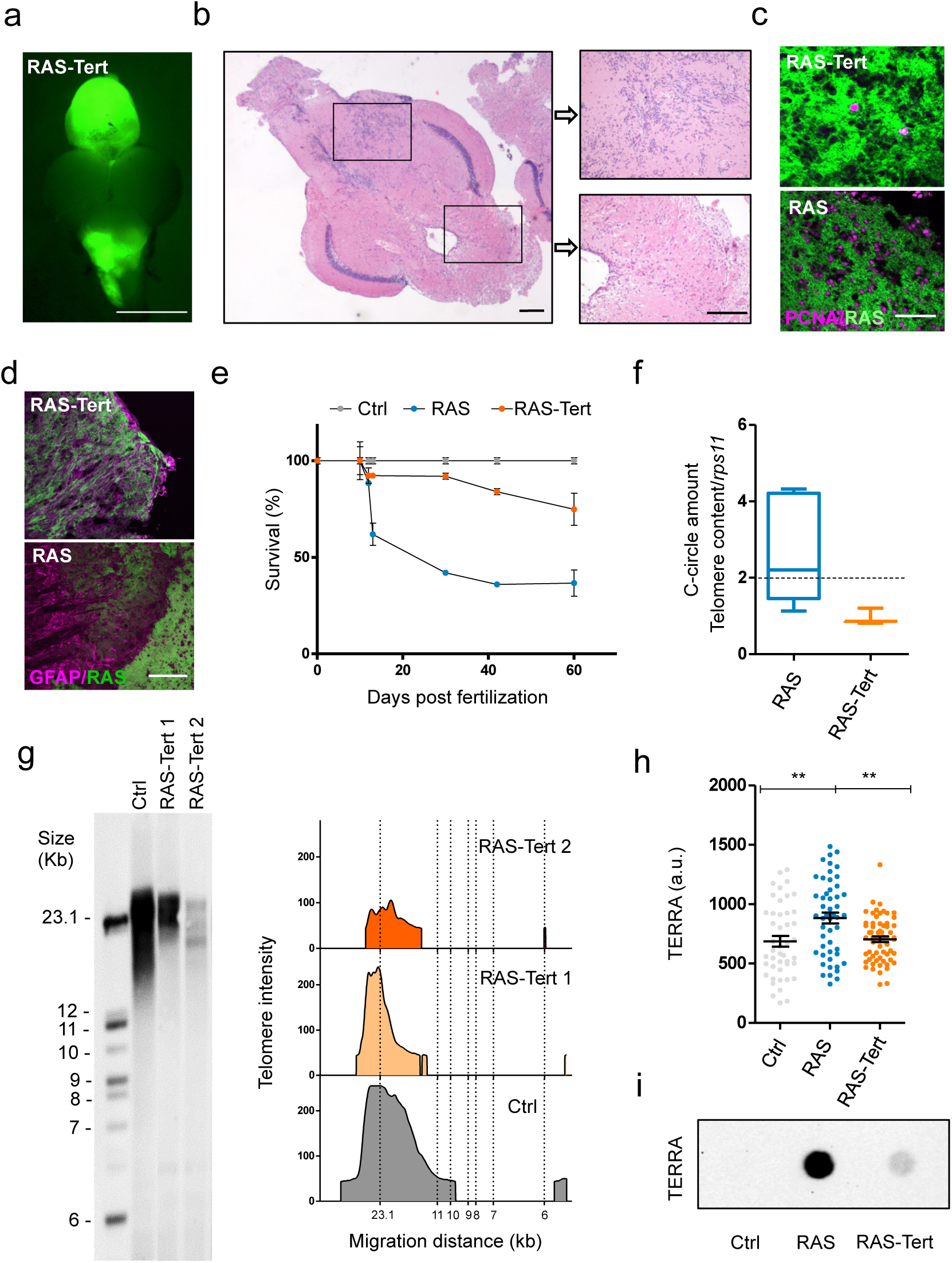
An overexpression of functional telomerase rescue ALT development. **a)** Representative fluorescent image of a RAS-Tert brain tumor. Scale bar 0.5 mm. **b)** Histological analysis of the RAS-Tert brain tumor shown in **a**, **c)** magnification of two area showing mild neoplastic abnormalities. Scale bars: 20 µm **c)** Immunofluorescence images showing the distribution of the proliferation marker PCNA (magenta) in sections of tumours from RAS-Tert and RAS brains. Scale bar: 20 µm **d)** Immunofluorescence images showing the differentiation marker GFAP (magenta) in sections of tumour of RAS-Tert and RAS brains. Scale bar: 20 µm **e)** Survival curve during the entire larval period up to 2 m of Control, RAS and RAS-Tert fish (n=45-60 larvae/genotype in three independent experiments). **f)** C-Circle assay quantified by telomere qPCR in RAS and RAS-Tert tumors. Data are represented as C-Circle amount normalized to telomere content (TC) and single copy gene (*rsp11*). RAS n=7; RAS-Tert n=3. The dashed line indicates the level above which ALT activity is considered significant. Whiskers box plots represent median: min to max values. **g)** Telomere length analysis by TRF in one control and two RAS-Tert tumors. The panel on the right shows TFR analysis obtained by graphing intensity versus DNA migration. **h)** Scatter plot of TERRA RNA-FISH signals measured as fluorescent intensity of single spots per nucleus in Control (n=46), RAS (n=50) and RAS-Tert (n=62) cell nuclei from brains and brain tumors. Data from a single, representative experiment are shown (** p < 0.01). Bars represent mean +/-s.e.m. **i)** TERRA RNA dot blot from total RNA of control, RAS and RAS:Tert brains. 500ng of total RNA was spotted for all samples.

The overexpression of *tert* in the RAS-Tert line was verified through RT-qPCR using primers designed between the 12 and 13 exons of the *tert* transcript (Figure S4d) and within the 3’UTR for the endogenous *tert* levels (Figure S4e), to distinguish total *tert* from endogenous *tert* levels. Total *terc* levels were not found increased compared to control fish as detected by RT-qPCR (Figure S4f).

We observed that the formation of neoplastic lesions due to induction of oncogenic RAS (Figure 6b) occurred with the same frequency in RAS-Tert and in RAS fish, being 100% in both cases, and in similar locations. In particular, in both genetic backgrounds, brain tumours appeared at 3-4 weeks in the telencephalon, IV ventricle and diencephalon (Figure 6a). Even though both RAS and RAS-Tert fish developed tumors with a similar frequency and timing, the two models differed significantly in overall survival, with the RAS-Tert fish showing 83.97% survival at 2 months whereas only 36.9% RAS fish survived by the same time (Figure 6c). These findings suggest that tumors developed in RAS-Tert fish are less aggressive than tumors arising in RAS fish.

In accordance with this view, histological analysis of several RAS-Tert brain tumours showed that they were less proliferative (Figure 6c, upper panel) and more differentiated, expressing glial fibrillary acidic protein, GFAP (Figure 6d, upper panel) than those that develop in RAS fish (Figure 6c-d, lower panels).

Next, we evaluated the presence of C-Circles (Figure 6f) in RAS-Tert tumors and found that the CCA products were reduced to the levels detected in control brains. Telomere length of RAS-Tert tumours was studied using TRF. In this analysis, we found that telomere length was very similar to that of control brain cells and observed a substantial reduction in telomere length heterogeneity in RAS-Tert compared to RAS tumours (Figure 6g, compare with Figure 1a). The assay repeated with a longer telomeric probe shows a TRF signal in RAS-Tert brain tumors typical of telomerase+ cells (Figure S4g). Moreover, the levels of TERRA in RAS-Tert tumors were comparable to TERRA levels detected in non tumoral samples, by RNA-FISH and RNA dot-blot analyses (Figure 6h-i). These findings suggest that the overexpression of *tert* prevented ALT development and TERRA increase in the zebrafish brain tumors analysed.

### The genes involved in the activation of the pre-replicative complex may play a role in the switch between ALT and telomerase-dependent TMM in brain tumors

Having established that the overexpression of *tert* in zebrafish brain tumours prevented the development of ALT, and knowing that ALT occurs frequently in human juvenile brain tumors, we took advantage of the near-isogenic zebrafish models that develop brain tumors and that differ only for the expression of telomerase or the development of ALT. We decided to perform global analysis of gene expression in brains with ALT or telomerase positive tumours in the attempt to identify molecular drivers of ALT development.

We performed transcriptome analysis through RNA-sequencing (RNAseq) of three ALT and three telomerase positive (+) tumours (Figure S5a). Principal component analysis (PCA) showed two separate clusters corresponding to ALT and telomerase+ brain tumours (Figure S5b). By comparing gene expression among the two different tumour types, we identified a limited set of genes that were significantly deregulated (Figure S5c, red dots). The analysis of differentially expressed genes (DEG) using DESeq2^42^, showed 366 differentially expressed (DE) genes (Figure 7a, Table S1). To investigate the biological pathways altered between ALT and Telomerase+ brain tumours, we first identified the human orthologues of the zebrafish DE genes (RAS-Tert vs RAS) using Biomart^43^ and Beagle^44^ and then we performed Reactome pathways enrichment analysis considering the most enriched pathways, according to adjusted p-value (padj < 0.1) (Table S2). We found that eight out of thirteen enriched pathways were strictly related to cell cycle and DNA replication (Figure 7b, Table S2), suggesting that these pathways act synergistically in the same network. The most significant pathways included “Cell cycle – mitotic” and “Activation of the pre-replicative complex” (padj < 0.05) (Figure 7b, Table S2).

**Figure 7.**
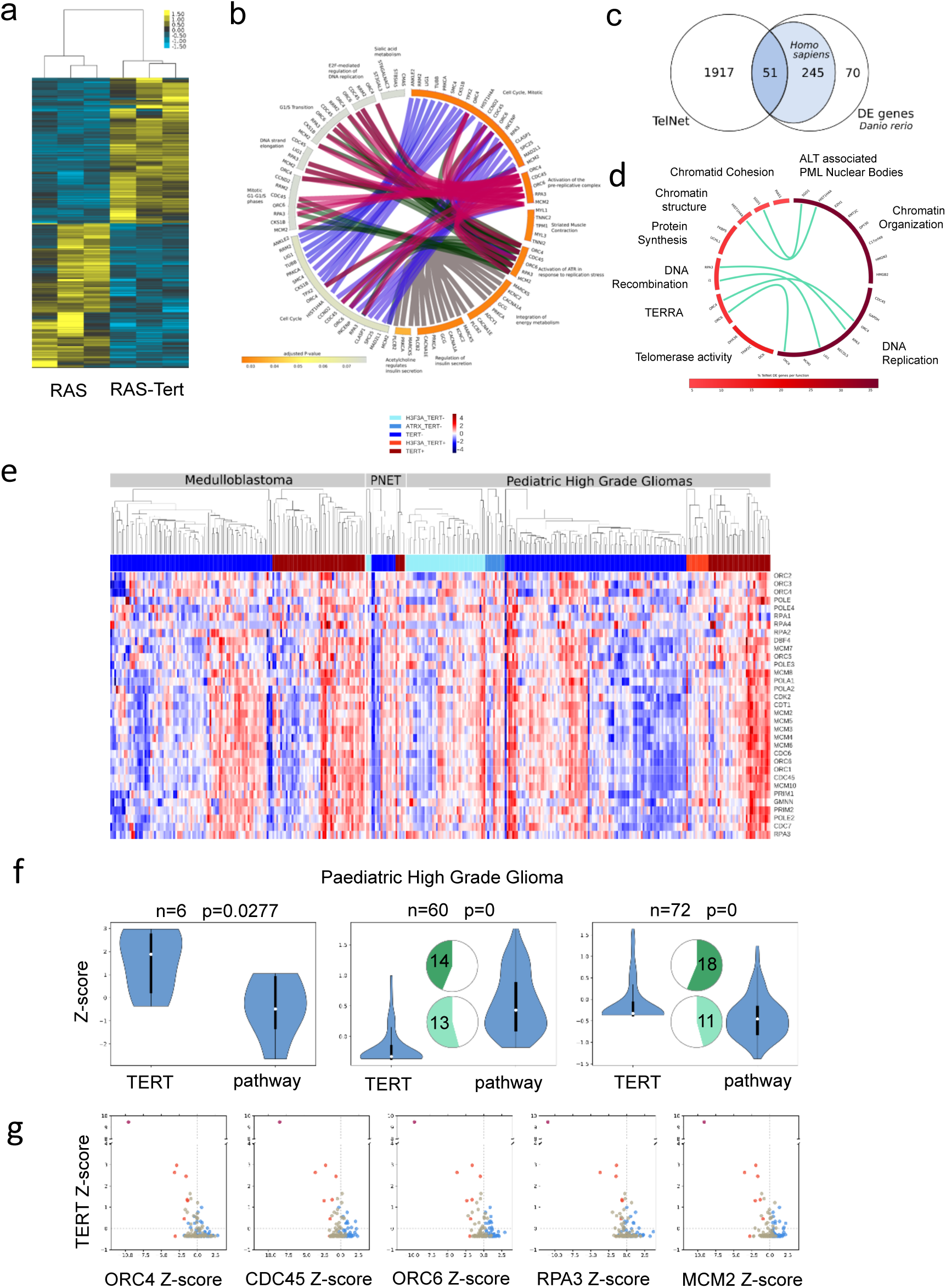
Analysis of RNA-Seq showed a class of genes altered in zebrafish brain tumours with different TMMs. **a)** Heatmap showing the 366 differentially expressed genes between brain tumors from RAS and RAS-Tert, hierarchically clustered with average linkage and uncentered correlation metric with cluster3, and displayed with tree view. **b)** Circular bar representing Reactome pathways enrichment for the DE gene set, colored according to adjusted P-value, with the most significant in orange, drawn with circos. DE genes in the pathways are indicated on the bars and are connected to the same genes in other pathways. **c)** Venn diagram showing the number of differentially expressed genes between RAS and RAS-Tert brain tumors, which have a human ortholog and are present in TelNet database. **d)** Circular plot depicting TelNet functions of the differentially expressed genes reported in TelNet database and their connections. **e)** Hierarchical clustering of the expression values of the genes of the Reactome pathway: “Activation of the pre-replicative complex” (Homo sapiens R-HSA-68962) in samples from the paediatric cBio Portal. The samples were divided per tumor type, according to the presence of mutations in H3F3A and/or ATRX, and TERT expression level (TERT+: Z-score >0, TERT-: Z-score <0). **f)** Violin plots comparing the expression of TERT and the mean expression of genes in the: “Activation of the pre-replicative complex” pathway in paediatric High Grade Glioma (pHGG). Samples for pHGG, were divided into three groups by K-means clustering. Pie chart plots show the percentage of samples (actual numbers are indicated) with H3F3A (dark green) or ATRX (light green) mutations for each of the three groups. **g)** Scatter plots of pHGG showing the expression of TERT (y-axis, in Z-score) vs the expression of five genes (*ORC4, CDC45, ORC6, RPA3,* and *MCM2*) in the: “Activation of the pre-replicative complex” pathway that were differentially expressed in zebrafish (x-axis, in Z-score). Samples marked in different colors belong to the three different groups as clustered by K-means.

We then evaluated whether the 366 DE gene cohort was enriched for genes known to regulate telomere biology. For this purpose, we used the Telnet database^45^ (http://www.cancertelsys.org/telnet/) which contains a list of genes that have been reported to be involved in telomere maintenance mechanisms (TMMs).

We established that the 17.3% of DE genes (51/296 DE genes with human orthologous) were listed in Telnet database (Figure 7c); we used the information provided by TelNet on gene-specific functions to classify the DEG identified in our brain tumour models. We found that the most significant DE genes belonged to the category: DNA replication and chromatin organization, according to the Telnet database (Figure 7d and Table S3).

Thus, Reactome pathway enrichment analysis and Telnet classification suggest that DE genes regulating DNA replication processes, especially the activation of the pre-replicative complex, plays a crucial role in the switch between ALT and telomerase in telomere maintenance in zebrafish brain tumors.

Therefore, we selected this pathway for further analysis in human data. We analyzed the expression of the 33 genes reported by Reactome (*Homo sapiens,* R-HAS-68962) as composing the “Activation of the pre-replicative complex” pathway (list of the genes in Table S4). We used the TCGA expression profiles of different paediatric/juvenile brain tumors (https://pedcbioportal.org/), including 133 medulloblastomas (MB), 20 primitive neuroectodermal tumors (PNET) and 138 pediatric high grade glioma (HGG) (Figure 7e, S6a-b). Moreover, we correlated the expression of the 33 genes involved in the activation of the pre-replicative complex pathway with *TERT* expression levels. Since no information was present about the TMM phenotype of these tumors, we also reported the presence of somatic mutations in the genes encoding H3.3 and ATRX, as these mutations are frequently found in ALT paediatric brain tumors^46^ (Figure 7e).

The hierarchical clustering on each tumor type (Figure 7e) suggests that all the genes of the pathway work as a module, when the pathway is activated, all the genes are upregulated and vice-versa, leading to the identification of different segregation groups. To understand if the pathway is activated in relation to low *TERT* expression, we clustered the data using K-Means (Figure 7f, S6c, e). We found three different groups, and the behaviour of *TERT* was anti-correlated with the mean expression of the pre-replicative complex genes in two out of three k-groups (Figure 7f, S6c, e). Mutations in H3F3A (pHGG n= 32; PNET n=3) and ATRX (pHGG n=24, PNET n=1) reported for some samples were clustered in group 2 (low expression of *TERT*, high expression of the genes of the pathway) and group 3 (*TERT* Z-score close to 0, pathway Z-score close to 0) (Figure 7f, S6e). However this analysis was not informative for all samples and we could not assign an ALT status to these or other samples based on gene expression only.

The profile of the 5 genes (*ORC4, CDC45, ORC6, RPA3* and *MCM2*) found deregulated in zebrafish RAS-TERT *vs* RAS tumors confirmed the presence of three different types of relationships: some tumors showed that when TERT was up-regulated (Figure 7g, S6 d,f, Y-axis Z-score >0) the five genes were down-regulated (x-axis Z-score<0) and vice-versa; the third group of samples showed that when the levels of TERT were close to zero, also the five genes of the pathway were close to zero (Figure 7g, S6 d,f).

Thus, both in zebrafish brain tumor models and in human juvenile brain tumors, the expression of telomerase is mostly anti-correlated with genes of the pre-replication complex, suggesting that in ALT tumors more active DNA replication may contribute to telomeric dysfunction.

### Identification of TMMs in a panel of twenty human juvenile brain tumours

To search for a definitive correlation between ALT and DNA replication in human brain tumors, we evaluated TMM in a panel of 20 primary, mostly juvenile, brain tumours of different histology. We performed the characterization of TMM in paraffin-embedded brain tumors, distributed as follows: four MB, five Central Nervous System Primitive NeuroEctodermal Tumors (CNS-PNET), one oligodendroglioma (ODG), one astrocytoma (AC), two juvenile glioblastomas (GBMs) and four rare histological variants of conventional GBM with Primitive Neuronal Component (GBM-PNC); three adult GBMs were added as controls (Figure 8a, Table S5).

**Figure 8.**
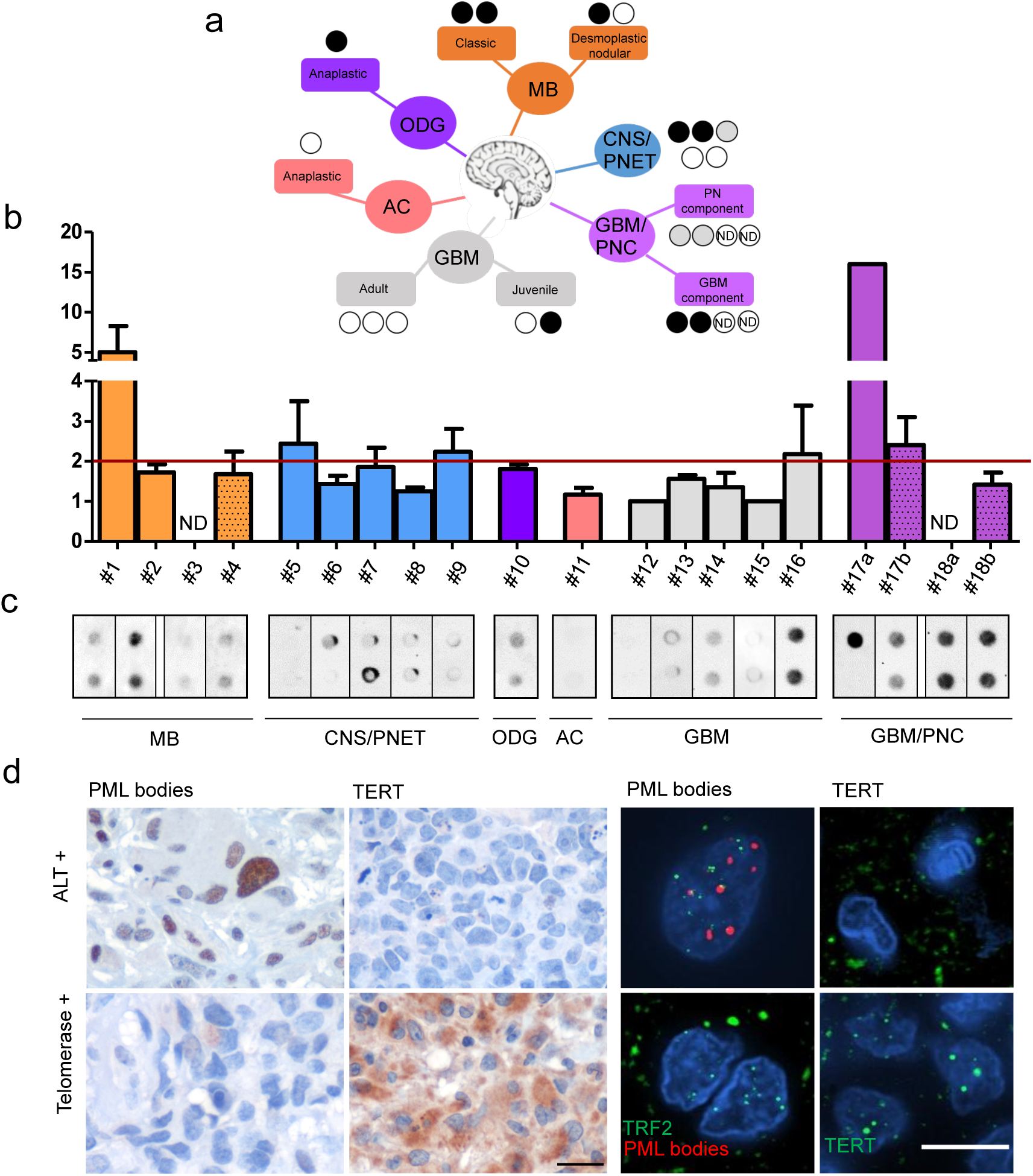
Identification of TMMs in a cohort of twenty human pediatric brain tumour. **a)** Tree representation of the human cases analysed here and the TMMs identified. Black, grey and white circles indicates the results of CCA analysis (black: positive, grey: almost positive and white: negative). **b)** Identification of human brain tumors positive or negative for C-Circles, analysed by qPCR (Bars represent mean +/-s.e.m of two independent experiments) and **c)** dot blot. The red line shows the limit above which ALT is detected. MB= Medulloblastoma, CNS/PNET =Central Nervous System Primitive NeuroEctodermal Tumors, ODG = oligodendroglioma, AC = astrocytoma, GBM = glioblastoma, GBM-PNC = Glioblastoma with Primitive Neuronal Component. **d)** Immunohistochemical and immunofluorescent stainings of PML bodies, TERT and PML bodies/ TRF2 in two representative cases of juvenile brain tumors, ALT or telomerase+, as indicated. Calibration bars: 10 µm

**Figure 9.**
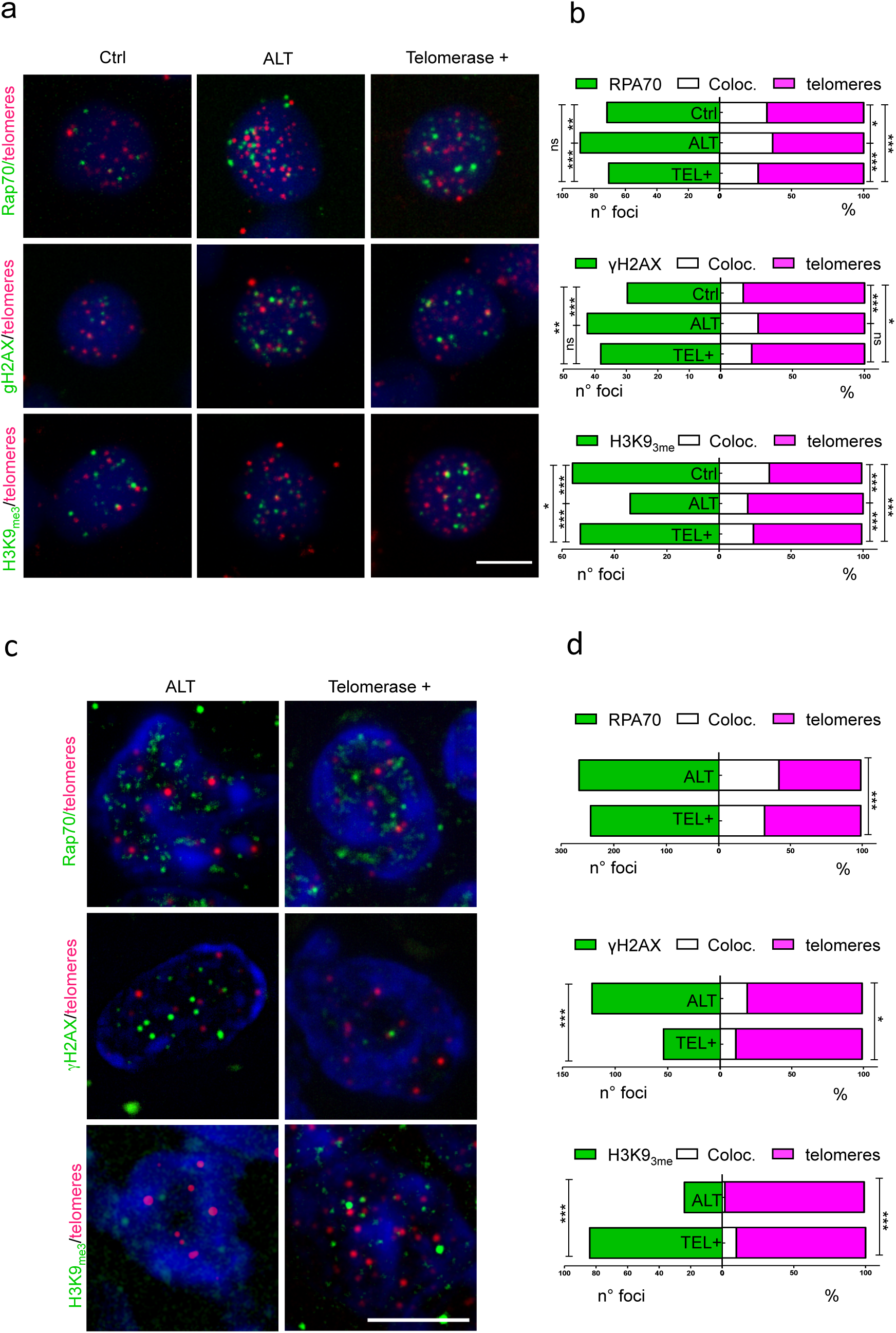
The presence of telomerase leads to a reorganization of telomeric chromatin, decrease DDR and replication stress. **a)** Fluorescent microscope images of representative control, ALT, and telomerase+ zebrafish brain tumor cells, stained via immunofluorescence (green) combined with QFISH (magenta). Antibody against replication stress marker (RPA70), DNA damage marker (γH2AX), and chromatin methylation marks (H3K9me3) were used and counterstained with DAPI. Scale bar: 5 µm **b)** Immunofluorescence quantification expressed as the number of foci per nucleus (green), and percent of immunofluorescence foci (white) that colocalized at telomeres (magenta) per nucleus (n=25-60 nuclei); * p < 0.05, ***p < 0.001 between the indicated groups. **c)** Fluorescent microscope images of representative ALT, and telomerase+ human brain tumor cells, stained via immunofluorescence (green) combined with Q-FISH (magenta). Antibody against replication stress marker (RPA70), DNA damage marker (γH2AX), and chromatin methylation marks (H3K9me3) were used and counterstaining with DAPI. Scale bar: 5µm. **d)** Immunofluorescence quantification expressed as the number of foci per nucleus (green), and percent of immunofluorescence foci (white) that colocalized at telomeres (magenta) per nucleus (n=25-60 nuclei); * p < 0.05, ***p < 0.001 between the indicated groups

First, we examined the tumours for the presence of C-Circles with Q-PCR and dot blot analysis (Figure 8b-c, Table S5). Using two methods for detecting C-Circles allowed us to identify ten ALT cases, including 3 MB (Table S5, no.1,2,4), 3 CNS-PNET (Table S5, no. 6,7,9), 1 ODG (Table S5, no.10), 1 juvenile GBM (Table S5, no.16) and 2 GBM/PNC (Table S5, no. 17,18) (black dots in Figure 8a, green and dark pink highlights in Table S5).

Generally, a positive C-Circle signal is considered a reliable indicator of ALT^35^, but we worried that the mostly single-stranded C-Circle signal could be lost in paraffin-embedded material during sample storage, while negative signals must be interpreted with caution. For this reason, we also evaluated the presence of PML bodies and the expression of TERT by immunohistochemistry (Figure 8d), for additional evidence supporting the activation of ALT in the positive cases (Table S5).

The ten C-Circle positive cases, were also positive for PML immunoreactivity, where PML foci co-localized with TRF2 (Figure 8d, upper panel; Table S5) and mostly negative for TERT (7 out of 10, Figure 8d, green highlight in Table S5) suggesting that ALT is the main TMM active in these tumours; three C-Circle/PML positive tumors, including 1 juvenile GBM (Table S5, no 16) and 2 GBM/PNC (Table S5, no.17,18) were also positive for TERT expression (dark pink highlight in Table S5).

Five tumours negative for C-Circles and PML bodies (1 out of 4 MB, 1 out of 2 juvenile GBM, 3/3 adult GBM, light blue highlight in Table S5) resulted positive for TERT (Figure 8d), suggesting that telomerase is the most representative TMM in these tumors.

For some cases, it was difficult to attribute a TMM classification. These included 1 out of 5 CNS-PNET tumours (no. 8 in Table S5), which showed low score for PML bodies and no signal for TERT and C-Circle, whereas 1 out of 5 PNET (no. 5 in Table S5) and 1 AC (no.11 in Table S5) resulted negative for all three markers.

A surprising result was relative to GBM/PNC tumours, where the presence of C-Circle, PML bodies and TERT was confined mostly to the GBM component, whereas the PNC component showed an absence of TERT expression and no PML bodies immunoreactivity (pink highlight in Table S5).

The molecular profile showed that only one case of the 10 ALT tumours presented the IDH1R132 mutation and the loss of ATRX (no.17 in Table S5).

This analysis allowed us to identify: a set of juvenile brain tumour which were clearly ALT (Table S5, green highlight), a set of brain tumours which appeared telomerase-dependent (Table S5, light blu highlight), a set of ALT/TERT+ tumors (Table S5, pink highlight), and a set of tumors with uncharacterized TMM mechanisms. These tumors with identified TMMs were then used for the analyses showed in the next set of experiments, aimed at localizing markers of replication/heterochromatin dysfunctions at telomeres of ALT or telomerase+ tumors.

In conclusion, ALT features frequently occur in different types of juvenile brain tumors, and the TMM status can be diagnosed from conventional pathology material.

### The reactivation of telomerase rescues ALT by preventing stalled replication forks and by reorganizing telomeric chromatin

Our analysis of human paedriatric brain cancers and zebrafish brain tumors revealed a reduced expression of genes of the pre-replicative complex and increased expression of genes driving heterochromatin deposition as hallmarks of the telomerase rescue. For this reason, we decided to search for signs of replication stress/stalled replication fork (RPA complexes), associated to increased double strand breaks (γH2AX foci), and assessed the status of heterochromatin (H3K9_me3_ foci) specifically at telomeres of ALT or telomerase+ tumor cells.

In both zebrafish and human tumours, we found a significant increase of telomeric RPA+ foci in ALT brain tumor cells compared to telomerase+ tumors (Figure S7a, c, upper panels). The number of telomeric RPA foci in telomerase positive zebrafish brain tumors was not significantly different from the number of RPA foci in zebrafish control brains, while it was increased in ALT tumors, suggesting that telomerase activity was able to reduce replication stress to the basal state.

Next, we investigated the presence of telomere dysfunction-induced foci (TIF, an indication of DNA damage) by immunostaining for γH2AX (Figure S7a, c, lower panels). Although there was no significant difference between γH2AX telomeric foci between ALT and telomerase+ zebrafish tumors, a difference was evident in human tumors, where the number and the localization of DNA damage foci at telomeres of ALT tumors were significantly higher than in telomerase+ tumors. This latter result demonstrates that in ALT human brain tumors the levels of DNA damage at telomeres is higher than in telomerase+ human brain tumors, suggesting that telomerase can resolve the replication stress, and reduce DNA damage, even if similar DNA damage levels may be present in non telomeric regions.

Finally, we investigated the distribution of heterochromatin foci at telomeres, using H3K9_me3_ as marker. We found a dramatic decrease in telomeric chromatin histone 3 lysine 9 methylation both in zebrafish and human ALT brain tumors (Figure 8e-f).

Thus, quantification of RPA, γH2AX and H3K9_me3_ foci localized at telomere in ALT or telomerase+ brain tumors suggests that the rescue of ALT by telomerase overexpression may be mediated by a reduction of telomeric DNA replication, with consequently reduced replication stress, and re-establishment of protective telomeric heterochromatin, which in human tumors also leads to a reduction of TIFs. As the effects of telomerase overexpression at the molecular level involve a reduction in the expression of genes of the pre-replicative complex, and a consolidation of genes involved in heterochromatin maintenance, it is tempting to speculate that telomere maintenance mechanisms act as major drivers of the coordination between DNA replication and chromatin status at telomeres in brain cancers.

## DISCUSSION

During recent years, the zebrafish has been used to study telomere biology in relation to organismal aging using mostly the telomerase mutant *hu3430*^31, 32, 37, 47^. Here, we characterize the telomere maintenance mechanism used by tumor cells in a zebrafish model of brain cancer, which highly resemble human paediatric glioblastoma of mesenchymal origin, a very aggressive tumor associated with poor prognosis. We found that zebrafish tumors rely on ALT as telomere maintenance mechanism, similarly to paediatric GBMs. ALT is thought to be triggered by factors that alter chromatin structure at telomeres; these include reduced deposition of histone 3 variants, due to defective histone chaperon ATRX activity or direct histone 3 mutations, altering H3K27 methylation, and heterochromatin formation. But how altered chromatin organization triggers ALT and whether the choice of TMM is a cause or a consequence of the telomeric chromatin status in cancer is not fully understood. By investigating the RAS zebrafish cancer model we report that ALT can develop not as a consequence of one of the mutations that alter chromatin structure at telomeric regions, but as a direct consequence of the lack of telomerase expression. Indeed, sustained expression of telomerase in the neural progenitors that initiate tumorigenesis prevented ALT development. Previous evidences in human cancer cell lines have shown that expression of hTERT does not abolish ALT^48^. However, genetic ablation of telomerase in a mouse cancer model can lead to ALT development^4^, suggesting that telomerase activity can prevent the emergence of ALT in cancer. In the zebrafish brain cancer model, sustained telomerase expression in tumor initiating cells not only prevents ALT, but also re-establishes the heterochromatin structure at telomeres, reduces DNA replication stress and represses TERRA transcription. Previous studies have indicated the shelterin complex as instrumental in organizing heterochromatin at telomeres and in preventing DNA damage in HeLa cells^49, 50^. On the basis of our observations, it would be interesting to know whether telomerase may be upstream of the shelterin complex in heterochromatin organization.

What is the role of telomeric DNA replication in ALT+ cells? Our study on DEG between ALT and telomerase+ brain tumors indicates that the pre-replicative complex may play a role in ALT development or maintenance. Expression of the genes of the pre-replicative complex is anti-correlated with *TERT* expression both in human paediatric and in zebrafish juvenile brain tumors. Kurth and Gautier^51^ have shown that pre-replicative complexes assemble within telomeric DNA and can be converted into functional origins. Indeed, the majority of telomeric DNA is duplicated by conventional DNA replication, but telomerase may control the amount of DNA replication at telomeres by binding and sequestering TRF2, which has been shown to recruit origin of replication complexes (ORCs) through its TRFH dimerization domain^52^. Thus in the absence of telomerase, an excess of ORC and TRF2 contribute to increase telomeric DNA replication, with the excess telomeric DNA being useful for recombination and for ALT associated C-Circles.

In ALT cells increase of DNA replication is linked to decrease heterochromatin formation. The open chromatin status may lead to the incorporation of non canonical variant repeats, which alter the binding of the shelterin complex, thus reinforcing the loop between DNA-replication and telomere de-protection. In addition, disruption of telomeric chromatin environment results in higher levels of TERRA transcription. TERRA transcripts may participate in ALT induction by multiple mechanisms: through formation of DNA:RNA hybrids, or R-loops, which may promote homologous recombination among telomeres^23^; by interfering with ATRX functions, as described in mouse embryonic stem cells^53^; by impacting replication of telomeric DNA^22^.

Heterochromatin is a feature of telomeric repeats and it is known to spread to subtelomeric regions in all species, maintaining a “silent” chromatin environment, but even more importantly, contributing to the 3D organization of the genome of differentiated cells. Cancer is challenging this organization, not only because it imposes cell and DNA replication at increase rates, but also because it is characterized by decreased DNA methylation and euchromatinization^54^. These changes involve telomeric chromatin, which must be protected, because of its fundamental role in 3D genome organization^55^; telomerase may play this role, and this is why it is often re-expressed in cancer, no net increase of telomere length is reported when telomerase is overexpressed. The non canonical role of telomerase, hypothesized by Chan and Blackburn^56^ may well include organization and maintenance of heterochromatin, inhibition of DNA replication in the telomeric region, and perhaps resolution of DNA damage resulting from stalled replication forks; indeed, it has recently been shown that, in human (as in yeast), telomerase recruitment to telomeres, besides being regulated by the shelterin complex^58^ is also driven by the DNA damage sensor kinases, ATM and ATR^59^.

Why is ALT more frequent (and prognostically worst) in paediatric tumors? It is plausible that reactivation of telomerase, which is usually achieved in cancer through mutations of the TERT promoter that hinder a repressor binding site, may require time for selection of the right mutation; therefore a quicker, although dangerous, mechanism to maintain telomere length and allow cancer growth, is through the engagement of ALT mechanisms. In addition, young cells, such those in developing brains or bones (for sarcomas), have probably not yet completely organized their heterochromatin, so that their hypomethylated chromatin environment makes it easier to switch to ALT. The mechanism is similar in the frequent mutations in H3.3 and ATRX in paediatric brain cancer, which compromise chromatin structure and allows for genomic instability and ALT to occur.

The use of a juvenile zebrafish model of brain cancer that progressively develops ALT allowed us to uncover a strong connection between an “immature” heterochromatin status and ALT development. In this scenario, the increase of telomeric DNA replication is a direct consequence of the lack of heterochromatin. It remains to clarify how is telomerase repressing the expression of genes of the pre-replicative complex and increasing the expression of those involved in heterochromatin organization. Very likely it is an indirect effect, although we cannot rule out that *tert* or *terc* may act directly on gene promoters to regulate their expression.

## METHODS

### Maintenance of zebrafish and line generation

Adult zebrafish (*Danio rerio*) were housed in the Model Organism Facility - Center for Integrative Biology (CIBIO) University of Trento and maintained under standard conditions^60^. All zebrafish studies were performed according to European and Italian law, D.Lgs. 26/2014, authorization 148/2018-PR to M. C. Mione.

Fishes with somatic and germline plasmid expression were generated as described^30, 41^. The following zebrafish transgenic lines were used or generated in the course of this study:

*Et(zic4:Gal4TA4, UAS:mCherry)^hzm5^* called zic:Gal4^30^

*Et(kita:Gal4TA4, UAS:mCherry)^hzm1^* called Kita:Gal4^41^

*Tg(UAS:eGFP-HRAS_G12V)^io006^* called UAS:RAS^41^

*hu3430 (Tert-/-)*^27^

*tg(10xUAS:tert) this study*

*tg(10xUAS:terc) this study*

### Cell culture and cell lines

The U2OS, HeLa, L5178Y-S and L5178Y-R lymphocyte cell lines were cultured in Dulbecco’s modified Eagle’s medium (DMEM) supplemented with 10% (v/v) fetal bovine serum (FBS) in a humidified incubator at 37 °C with 5% CO2. Cell lines were tested regularly for mycoplasma contamination by Celltech CIBIO facility.

### Human pathology material

Human brain tumour samples were retrieved from the archive of Department of Pathology, Spedali Civili di Brescia, University of Brescia, in agreement with protocols approved by the Institutional Review Board. Specifically, for the retrospective and exclusively observational study on archival material obtained for diagnostic purpose, patient consent was not needed (Delibera del Garante n. 52 del 24/7/2008 and DL 193/2003).

### DNA constructs and transgenic line generation

The genes encoding zebrafish terc (ENSDARG00000042637.10) and tert (ENSDARG00000042637.10) were synthesized and cloned in pBluescript II KS+ and subcloned in a pEntry vector (pME-MCS Tol2Kit, http://tol2kit.genetics.utah.edu/). The UAS:terc; cmlc2:eGFP and UAS:tert; cry:eGFP constructs were generated by MultiSite Gateway assemblies using LR Clonase II Plus (Life Technologies) according to standard protocols and Tol2kit vectors described previously^61^. 25 pg of the final construct and 25pg of tol2 transposase mRNA were injected into 1-cell stage embryos and founder fish (F0) for terc or tert were identified based on green fluorescent heart or fluorescent eyes. Embryonic brain expression was obtained by outcrossing them with the zic:Gal4 line.

### Terminal restriction-fragment (TRF)

Telomere length assay was performed as described^62^. Briefly, genomic DNA was extracted from freshly isolated zebrafish brains following the instruction of ReliaPrep™ gDNA Tissue Miniprep System (Promega Corporation) and then 3µg of genomic DNA were digested with RsaI and HinfI enzymes (New England Biolabs) for 12 h at 37°C. After electrophoresis, the gel was blotted and probed with antisense telomere probe (CCCTAA)_8_ labelled with DIG Oligonucleotide 3’-End labeling Kit (Sigma-Aldrich) or with a 1.6 kb fragment containing the sequence (TTAGGG)n^62, 63^ labelled with Nick Translation Kit (Sigma-Aldrich). Image Lab™ Software (Biorad) was used to analyse telomere length from TRF analysis, data were plotted using GraphPad Prism.

### Quantitative fluorescence in-situ hybridization (Q-FISH) on interphase nuclei

Q-FISH on interphase nuclei was performed as described^32^. Cell suspensions were obtained as described^32^. Z stack images were acquired with an inverted Leica DMi8 fluorescent microscope equipped with a monochromatic Andor Zyla 4.2 Megapixel sCMOS camera, using an HC PL Apo CS2 63x/1.4 oil immersion objective. Z-stack images were processed to remove background using Fiji/ImageJ and then telomere fluorescence signals were quantified using the TFL-TELO program (from Peter Lansdorp, Vancouver, Canada). Data were plotted using GraphPad Prism.

### Telomerase activity assay

Real-time quantitative TRAP (Q-TRAP) assay was performed as described^27^. Protein extracts were obtained from brain (Ctrl and *tert-/-*) and brain tumors (RAS). The *hu3430 tert-/-* mutant strain was used here. After PCR, real-time data were collected and converted into Relative Telomerase Activity (RTA) units based on the following formula: RTA of sample =Delta10^(Ct^ ^sample-Yint)^/slope.

### Analysis of Gene Expression

Total RNA was extracted from larval heads and brains/tumors with TRIzol reagent (Invitrogen). Total RNA was cleaned up using RNeasy Mini Kit (Qiagen) following the manufacturer’s instructions and treated twice with DNase I (1 unit/µg RNA, Qiagen). The RNA concentration was quantified using nanodrop2000 (Thermo Fisher) and VILO superscript KIT (Thermo Fisher) was used for First-strand cDNA synthesis according to the manufacturer’s protocol. qRT-PCR analysis was performed using qPCR BIO Sygreen Mix (Resnova - PCR Biosystem) using a standard amplification protocol. The primers used for *zebrafish tert* were: forward 5’-CGGTATGACGGCCTATCACT-3’ and reverse 5’-TAAACGGCCTCCACAGAGTT-3’; for 3’ UTR *zebrafish tert* were: forward 5’-AACACTTGATGGTGACTGT-3’ and reverse 5’-GACTTCTGCATCGATCTGTGAT-3’; for *zebrafish rps11*: forward: 5’-ACAGAAATGCCCCTTCACTG-3’ and reverse: 5’-GCCTCTTCTCAAAACGGTTG-3’; for *human* 36B4: forward 5’-CAGCAAGTGGGAAGGTGTAATCC-3’ and reverse: 5’-CCCATTCTATCATCAACGGGTACAA-3’. To determine *terc* and TERRA levels, total RNA (1 µg) was reverse transcribed with gene-specific primers (zebrafish *terc:* 5′-TGCAGGATCAGTGTTTGAGG-3’; *rps11*: 5’ -GCCTCTTCTCAAAACGGTTG-3’; Telo RT 5′-CCCTAACCCTAACCCTAACCCTAACCC TAA-3′, *human* 36B4 RT 5′-CCCATTCTATCATCAACGGGTACAA-3′) using Superscript III (Thermo Fisher) at 55 °C for 1 h, followed by RNase H treatment. qRT-PCR analysis was performed using qPCR BIO Sygreen MIx (Resnova - PCR Biosystem) with 500 nM specific primers (*zebrafish terc* forward: 5′-GGTCTCACAGGTTTGGCTGT3′, reverse 5′-TGCAGGATCAGTGTTTGAGG-3′); (*zebrafish TERRA* forward: 5′-CGG TTT GTT TGG GTT TGG GTT TGG GTT TGG GTT TGG GGT-3′, reverse 5′-GGC TTG CCT TAC CCT TAC CCT TAC CCT TAC CCT TACCCT-3′). *rps11* and *36B4* specific primers were used as zebrafish and human reference. The amplification program was as follows: 95°C for 10 min followed by 36 cycles at 95°C, 58°C and 72°C each for 10 s. Real-time PCR was performed with a CFX96 Real-Time PCR Detection System (Bio-Rad) machine. Q-PCR analysis was performed with Microsoft Excel and Graphpad Prism. In all cases, each PCR was performed with triplicate samples and repeated with at least two independent samples. Reactions without reverse transcriptase were performed as controls for TERRA quantification.

### Zebrafish 5mC tert promoter ChIP protocol

Larval heads and brains/tumors samples were incubated for 8 minutes at room temperature in 500µl 1% formaldehyde in PBS + protease inhibitors. Then 50µl of Glycine 1.25M were added to stop cross-linking reaction, samples were briefly vortexed and incubated for 5 minutes at room temperature on a wheel. Upon centrifugation, pellets were washed in 1.2 ml ice-cold PBS, resuspended in 500 µl of PBS and homogenized using a homogenizer. After homogenization, samples were centrifuged 5000 rpm for 10 minutes at 4°C and pellets resuspended in 600 µl of Lysis buffer (50mM Tris pH8; 10mM EDTA pH8; 1% SDS) + protease inhibitors and incubated 10 minutes in ice. Samples were vortexed for 30 seconds then centrifuged 1000rpm for 1 minute at 4 °C. The supernatant was collected into a new tube while the pellet was resuspended in 300 µl of Lysis buffer + protease inhibitors. After vortex and spin, as before, the supernatant was collected and added to the previous 600 µl of sample making a 900 µl lysate for each sample. The lysate was then divided in three tubes and sonication performed using the Bioruptor instrument (DIAGENODE) with the following setting: 4 x cycles 20 seconds ON and 60 seconds OFF at 4 °C. After sonication, 8 µl of lysate are run on agarose gel to verify chromatin shearing. 300-600nt long DNA fragments are expected. Sonicated samples were centrifuged 5000 rpm for 5 minutes at 4°C. The supernatant was collected in a new tube. At least 100 µg of proteins were used for subsequent IP. One-tenth of the amount of proteins used for IP is collected for the INPUT and frozen at -80 °C. 3.6 ml of dilution buffer (20mM Tris pH8; 150 mM NaCl; 2mM EDTA pH 8; 1% Triton) were added to each sample to be immunoprecipitated. A pre-clearing step was performed by adding 40 µl of protein G Dynabeads for 2 hours at 4 °C on a wheel. Upon Dynabeads removal each sample was then divided into two new tubes, in which 2 µg of either anti-5mC (Abcam) or IgG (Abcam) were added. Antibody incubation was performed overnight at 4 °C on a wheel. The following morning, 20 µl of pre-equilibrated protein G-Dynabeads were added to each IP and samples were incubated at 4 °C for 2 hours on a wheel. For each sample, Dynabeads-chromatin complexes were recovered using magnetic rack and washed three times with 1 ml wash buffer and one time with the final wash buffer. Each wash was performed for 5 minutes at room temperature. Then 450 µl of elution buffer (1% SDS; 0.2M NaCl; 0.1M NaHCO3) + 18 µl of NaCl 5M solution + 5µl of RNAse A solution (10mg/ml solution) were added to each sample. Samples were vortexed vigorously and incubated at 37 °C for 1 hour. Then 250 µg of proteinase K was added to each sample and incubated at 65 °C overnight. The following morning DNA was extracted using phenol/chloroform protocol, precipitated with Ethanol 100%-Sodium-Acetate solution in the presence of 20 µg glycogen, washed in Ethanol 70%, air-dried and resuspended in 30 ul (IP) water. Every step was performed also on the INPUT.

### C-Circles assay

C-Circles assay was performed as described^35^. Briefly 30 ng of genomic DNA was combined with 0.2 mg/ml BSA, 0.1% (v/v) Tween20, 1mM each dATP, dTTP, dGTP, 4 µM dithiothreitol (DTT), 1x Φ29 DNA polymerase buffer, 7.5 U Φ29 (New England Biolabs). Rolling Circle Amplification (RCA) reaction was performed by incubation at 30°C for 8 h, plus 20 min at 65 °C. Reactions without the addition of Φ29 polymerase were included as a control (“–Φ29”). For dot blot detections, the CCA products (plus 40 µl 2x SSC) were dot-blotted onto 2x SSC-soaked positive nylon membrane, thanks to a 96-well Bio-Dot Microfiltration Apparatus (Biorad). The membrane was UV-crosslinked for 3 minutes/each side and hybridized with probe (CCCTAA)_8_ labelled with DIG Oligonucleotide 3’-End labeling Kit (Sigma-Aldrich) and developed as described^62^. Image Lab™ Software (Biorad) was used to analyse dot intensity. The result of the C-Circle assay dot blot was evaluated according to^35^. Q-PCR detection was performed as described^31^. Briefly, CCA products were diluted 4 times in water and used as templates in a qPCR using telomF (300 nM) 5’-GGTTTTTGAGGGTGAGGGTGAGGGTGAGGGTGAGGGT-3’ and telomR (400nM) 5’-TCCCGACTATCCCTATCCCTATCCCTATCCCTATCCCTA-3’ primers. qPCRs using *rps11* primers (150 nM) and 36B4 primers (Forward 300 nM and Reverse 500 nM) were performed for Single Copy Gene (SCG) normalization in zebrafish and human samples, respectively.

All qPCRs were done in triplicates. Each telomere Ct was normalized with the SCG Ct (normTEL). The telomere content (TC) of a sample was the normTEL value in the –Φ29 reactions. The CC abundance of a sample was calculated as (normTEL in +Φ29) - (normTEL in –Φ29). ALT activity was considered significant if at least twice than the levels without Φ29 polymerase^35^.

### TERRA dot-blot

Before blotting, 500ng of total RNA (in 1 mM EDTA, 50% formamide - Volume 100 µl) were denatured in a thermocycler at 65 °C for 10 minutes and then on ice. Denaturated RNA was dot-blotted onto 2x SSC-soaked positive nylon membrane and then UV crosslinker for 3 min/each side. Hybridization was performed at 50°C O/N with the probe 1.6 kb fragment containing the sequence (TTAGGG)n^63^ labelled with Nick Translation Kit (Sigma-Aldrich) and developed according to^62^. Image Lab™ Software (Biorad) was used to analyse dot intensity; quantitative analysis of dot blot intensity was performed after background subtraction and on control normalization. Data were plotted GraphPad Prism. A digoxigenin-labelled actin mRNA sense probe, obtained from *in vitro* transcription, was used as loading control.

### TERRA RNA-FISH

Cell-derived from zebrafish brains and tumor brains were seeded on poly-D Lysine (1µg/mL) (Sigma-Aldrich) slides 1 hour before starting the experiment. TERRA RNA–FISH assay was performed as described^40^. Z-stack images were acquired with an inverted Leica DMi8 fluorescent microscope equipped with a monochromatic Andor Zyla 4.2 Megapixel sCMOS camera, using an HC PL Apo CS2 63x/1.4 oil immersion objective. Z-stack images were processed for removing background using Fiji/ImageJ and fluorescence signals were quantified using the TFL-TELO program (from Peter Lansdorp, Vancouver, Canada). Data were plotted using GraphPad Prism.

### Two-color Chromosome Orientation FISH (CO-FISH) in metaphase spreads

Metaphase spreads were obtain from larval heads and brains/tumors from adult individuals. For larvae: 30hpf embryos, previously dechorionated, were incubated with BrdU 10mM//BrdC 4mM (Alfa Aesar™), 1% DMSO for 6h at 28°C and then with 1µg/µl Nocodazole (Sigma-Aldrich) for an additional 6h before preparing cell suspensions. For adults, 5µl of BrdU/C and Nocodazole (at the concentration reported above) were retro-orbital injected^64^ with 12 hours interval; fishes were processed 24h after the first injection. Cell suspensions were obtained as previously described. Cells were then incubated for 25 minutes at 28.5°C in a hypotonic solution (2.5g/L KCl, in 10mM HEPES, pH 7.4) and fixed in ice-cold Carnoy’s methanol: glacial acetic acid fixative (3:1, v:v) per 2h. After a wash in Carnoy’s methanol, cells were dropped onto superfrost microscope slides (pre-cleaned and wet) and allowed to dry overnight at room temperature in the dark. Degradation of newly synthesized strand and 2-Color FISH was performed as described^36^. Metaphase spread chromosomes were counterstained with DAPI. Z-stack images images were acquired with an inverted Nikon Ti2 fluorescent microscope equipped with a monochromatic Andor Zyla PLUS 4.2 Megapixel sCMOS camera, using a Plan Apochromatic 100x/1.45 oil immersion objective. Images were processed for background subtraction using Fiji/ImageJ

### Immunostaining on paraffin-embedded sections

Briefly, 2-µm-thick paraffin sections were deparaffinized and rehydrated. Endogenous peroxidase activity was blocked with 0.3% H_2_O_2_ in methanol for 20 minutes. Antigen retrieval (when necessary) was performed in either 1.0 mM EDTA buffer (pH 8.0) or 1 mM Citrate buffer (pH 6.0). Sections were then washed in TBS (pH 7.4) and incubated primary antibodies diluted in TBS 1% BSA at room temperature for 1 hour. The reaction was revealed by using Novolink Dako EnVision+Dual Link System Peroxidase (Dako Cytomation) followed by DAB and slides counterstained with Hematoxylin. For double immunohistochemistry, after completing the first immune reaction using the anti-PML antibody, the anti-TRF-2 was revealed by Mach 4 Universal AP Polymer kit (Biocare Medical) using Ferangi Blue (Biocare Medical) as chromogen and nuclei were counterstained with Methyl Green. For immunofluorescence, secondary antibodies conjugated with FITC and/or Texas Red was used and nuclei were counterstained with DAPI. The antibody used and their dilutions were as follows: PML bodies 1:100 (Santa Cruz Biotechnology, cat. No.SC-966); TERT 1:100 (Novusbio, cat. No.NB100-317); TRF-2 1:200 (Novusbio, cat. No. NB110-5713055); PCNA 1:200 (Santa Cruz Biotechnology, cat. No. SC-25280); GFAP 1:1000 (Dako, cat. No.20334). Images were acquired through an Olympus DP70 camera mounted on an Olympus Bx60 microscope using CellF imaging software (Soft Imaging System GmbH).

### Immunofluorescence combined with Q-FISH

Cell suspensions derived from zebrafish brain tumors were seeded on Poly-lysine (1µg/mL) (Sigma-Aldrich) slides, whereas paraffin sections from human brain tumor were deparaffinized and rehydrated as described above before processing for immune-fluorescence. Briefly, after 1 wash in TBS 1x per 5 minutes, slides were fixed in 2% paraformaldehyde containing 2% sucrose for 10 minutes at RT and then washed twice in TBS, followed by permeabilization with 0.5% Triton for 15 minutes. After 3 washes in TBS, slides were incubated 1 hour at RT in blocking buffer (5% NGS, 0.1% Triton for H3K9_me3_ or 0.5% BSA, 0.2% Gelatin cold water fish skin in 1x PBS) and then overnight at 4°C with primary antibody with the following dilutions: RPA70 1:200 (Invitrogen, cat. No.PAS-21976), γH2AX 1:300 (Millipore, cat. No05-636), H3K9_me3_ 1:500 (Abcam, cat no. ab8898). After three washes in blocking buffer, slides were incubated with secondary antibody (goat-anti-mouse 488 or Goat-anti-rabbit 488 -Thermo Fisher) 1:500 for 2h at RT. After incubation, slides were washed 3 times in 1xTBS (5 minute each) and 1 time in Q-FISH washing buffer (0.1%BSA, 70% formamide, 10 mM Tris pH 7.2). Then FISH was performed with PNA TelC-Cy3 probe (PANAGENE) as described^39^. Nuclei were counterstained with DAPI. Z-stacks Images were captured at 100x magnification (Plan Apochromatic 100x/1.45 oil immersion objective) using an inverted Nikon Ti2 fluorescent microscope equipped with a monochromatic Andor Zyla PLUS 4.2 Megapixel sCMOS camera. Images were processed for background subtraction using Fiji/ImageJ. Colocalization analysis was performed with DiAna (ImageJ - https://imagejdocu.tudor.lu/doku.php?id=plugin:analysis:distance_analysis_diana_2d_3d_:start)^65^, calculating co-localization between objects in 3D, after 3D spot segmentation.

### RNA-Sequencing analysis

Demultiplexed raw reads (fastq) generated from the Illumina HiSeq were checked using FASTQC tool (Version 0.11.3). All samples passed the quality standards. Then we aligned to the reference genome Danio rerio assembly GRCz11 using STAR^66^ with recommended options and thresholds (version 2.5) HTSeq-count^67^ (version 0.9.1) was used to generate raw gene counts. Counts normalization to Trimmed Mean of M-values (TMM) for visualization methods was performed by edgeR package^68, 69^ (v.3.24.3).

The differential expression analysis was performed using DESeq2 package^42^ (v.1.22.2) and for significance testing, the Wald test was used. Genes were considered differentially expressed with adjusted P- value <0.05 and a log2 fold change greater than 1 or smaller than -1. For statistical analyses, the adjusted p-values were generated via the Benjamini-Hochberg procedure. For visualization, differentially expressed genes (Trimmed Mean of M-values, TMM) were hierarchically clustered with average linkage and uncentered correlation metric with cluster3^70^ and displayed with treeview^71^. Human orthologs were identified through Beagle Database (accessed in January 2019, and BioMart^43^ for 295 of the 366 differentially expressed genes). Functional annotation on human orthologs of differentially expressed genes was carried out through enrichr web tool^72^ (accessed in May 2019) on Reactome pathways. Pathways were represented using Circos plots using Circos package^73^. Connection curves between pathways represent the same gene is present in both pathways, they were manually drawn. Genes involved in telomere maintenance were obtained from TelNet database^45^. Expression data (Z-scores) for the Activation of the pre-replicative complex (Homo sapiens R-HSA-68962) Reactome pathway and TERT, were downloaded from the Pediatric cBio Portal^74, 75^ for samples whose mutations and RNA Seq V2 data were both available, in the following cancer types: medulloblastoma (MB), primitive neuroectodermal tumor (PNET), pediatric high grade glioma (pHGG). Samples per tumor type were divided in groups according to TERT expression level (TERT+: Z-score >0, TERT-: Z-score <0) and the mutations identified (H3F3A, ATRX), and hierarchical clustering with euclidean metric was performed on each group. Per tumor type, the downloaded Pediatric cBio Portal expression data were clustered using K-Means. TERT expression was subtracted from the mean expression of the pathway genes, for each sample. The number of groups (k) was determined using the elbow method. The difference between the expression of TERT and the mean expression of the pathway was tested with the Wilcoxon signed-rank test.

### Data availability

The raw data have been deposited in Gene Expression Omnibus Database, accession no. GSE134135.

### Statistics

Except for data from RNA-Seq analysis, all the graph and the statistical analysis (Mann-Whitney - nonparametric test, no Gaussian distribution, two-tailed, interval of confidence: 95%) were generated and calculated using GraphPad Prism software version 5.0. For all experiments a minimum of three fish or groups per genotype was used, unless differently specified. Details regarding number of samples used and statistical analysis are provided in the figure legends.

## ACKNOWLEDGEMENTS

We thank members of Mione’s laboratory for discussion and technical help; Veronica Bergo and GianMarco Franceschini for initial bioinformatics analysis; Giorgina Scarduelli, Sara Leo and Michela Roccuzzo of the AICF – Cibio Dept, for providing assistance in image acquisition and analysis; Veronica De Sanctis and Paola Fassan of the NGS-Cibio Dept, for library preparation and RNA-Seq. We thank Diana Garcìa, Jesus Garcia Castillo and Francisca Alcaraz of Cayuela’s laboratory for initial sharing of protocols and expertise on the CCA and Q-TRAP techniques. A.I. received support from Fondazione Veronesi through a postdoctoral fellowship and from The Company of Biologists - Disease Models & Mechanisms for a Travelling Fellowship. E.C. was supported by a Rita Levi Montalcini fellowship from the Italian Ministry of Education, University and Research (MIUR). M.L.C. was supported by the Spanish Ministry of Science, Innovation and Universities (grants PI16/00038), Fundación Séneca-Murcia (grant 19400/PI/14) and Fundación Ramón Areces. M.C.M. was supported by World Wide Cancer Research, grant no. 0624, for the generation of the brain tumor model, and by LILT –Trento, Program 5 per mille (year 2014).

## AUTHOR CONTRIBUTIONS

A.I. and M.C.M. conceived the study and performed all the experiments and the analyses, with the exception of the *tert* promoter ChIP experiments, that were performed by E.C. Histology, immunohistochemistry and immunofluorescence stainings and analysis of human speciments were performed by F.P. and P.L.P.; the generation of the *tg(10xUAS:tert)* and *tg(10xUAS:terc)* zebrafish transgenic lines was performed by M.B. and M.L.C., and the optimization of the Q-FISH, 2-colour CO-FISH and metaphase preparations was guided by F.B. All bioinformatics analysis was done by E.K. and S.P. Writing was done by A.I., E.C. and M.C.M. All authors commented on the manuscript.

