## Supplementary figures 1-6 for "Expression of telomerase prevents ALT and maintains telomeric heterochromatin in juvenile brain tumors"

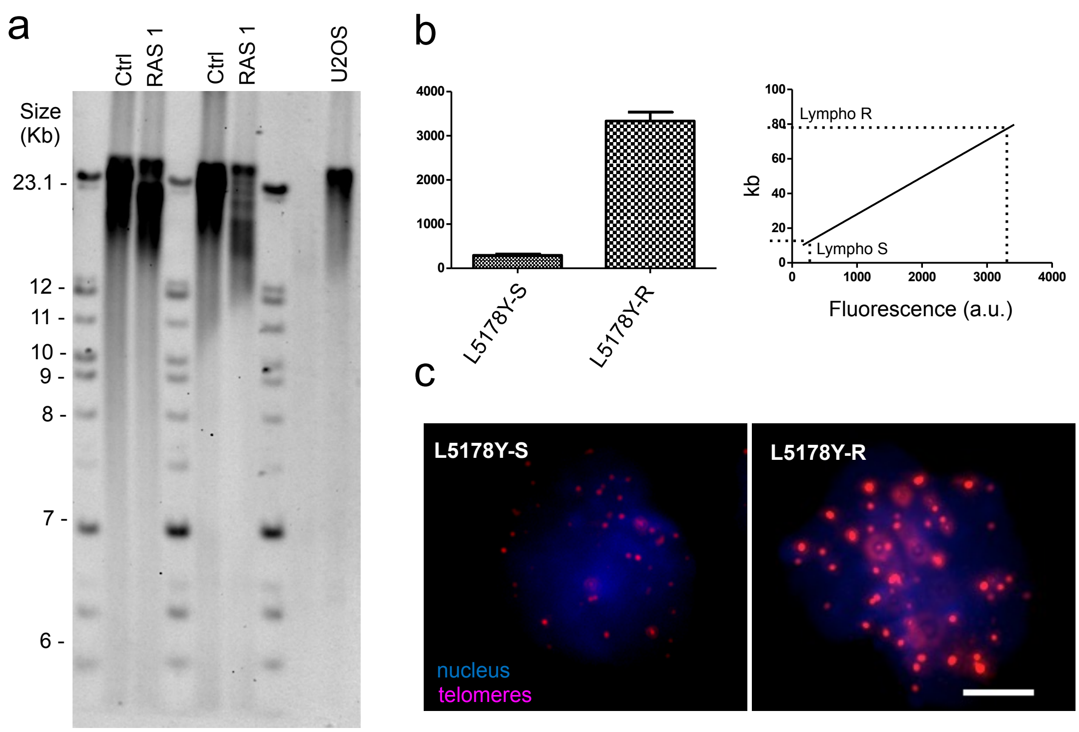


**Figure S1 (related to Figure 1: Zebrafish brain tumors have long and irregular telomeres)**

**a)** Terminal Restriction Fragment analysis of telomeres in Controlbrains and RAS tumors using a 1.6 kb DNA probe. Alt positive U2OS cells for comparison are in the last lane.
**b)** Representative panels describing how fluorescence intensity was correlated to length in kilobases (kb) using L5178Y-S and L5178Y-R lymphocyte cell lines with known telomere lengths of 10.2 and 79.7 kb respectively. The calibration was performed for each experiment.
**c)** Representative fluorescence images of Q-FISH performed in L5178Y-S and L5178Y-R lymphocyte cells, as indicated. Scale bar: 5μm.

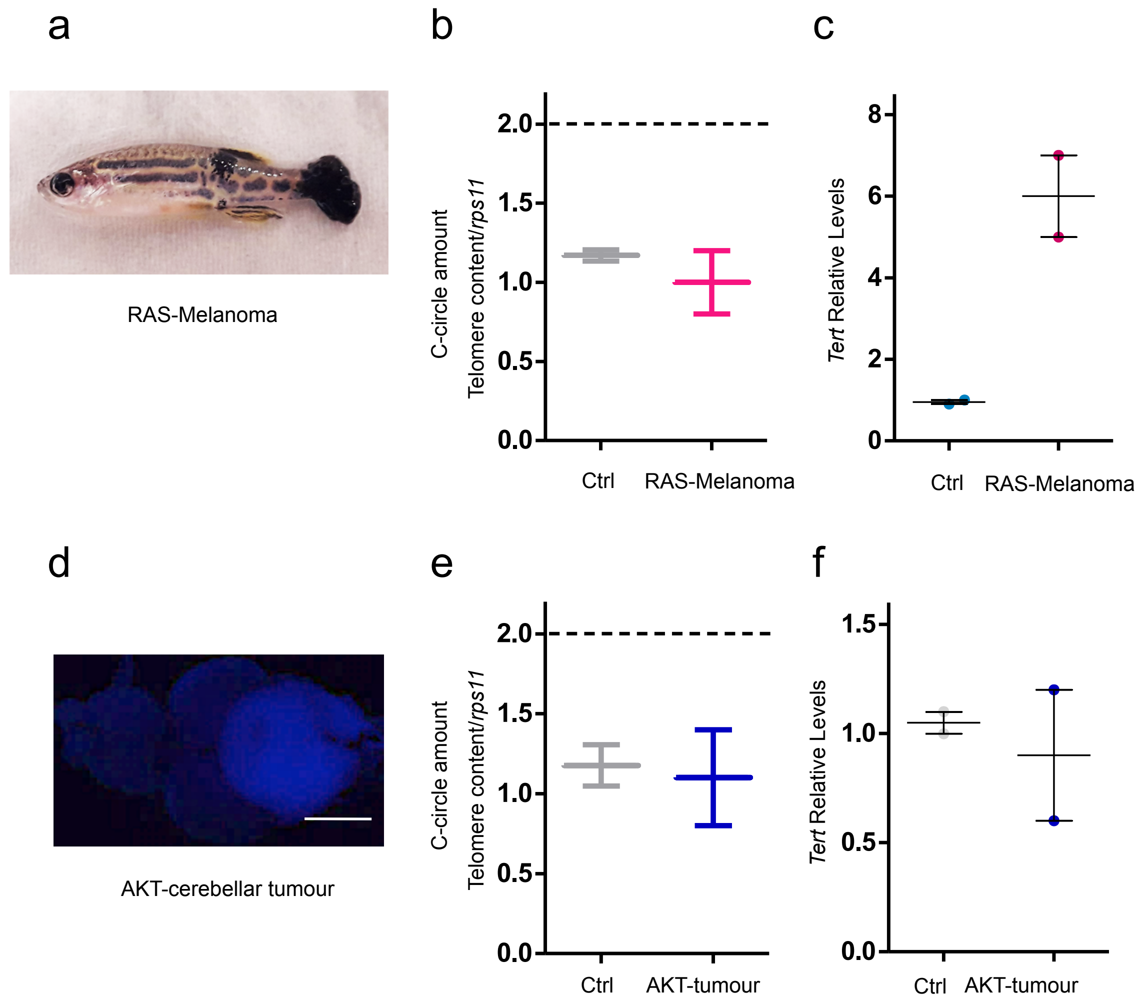


**Figure S2 (related to Figure 3: Zebrafish brain tumors are ALT-positive)**

**a)** Image of a fish with melanoma induced by overexpression of oncogenic HRAS$^{V12}$
**b)** C-Circle quantification by telomeric qPCR in control skin and melanoma (n=2).
**c)** Measurement of *tert* expression in control skin and melanoma. Values were normalized first to *rps11 mRNA* and then to *tert* expression in control skin (n =2).
**d)** Image of a zebrafish brain with an AKT - driven cerebellar tumor. Calibration bar: 0.5 mm
**e)** C-Circle quantification by telomeric Q-PCR in control brain and AKT - driven cerebellar tumors (n=2)
**f)** Measurement of *tert* expression in control brains and AKT - driven cerebellar tumors. Values were normalized first to *rps11* and then to *tert* expression in control brains (n=2)

a

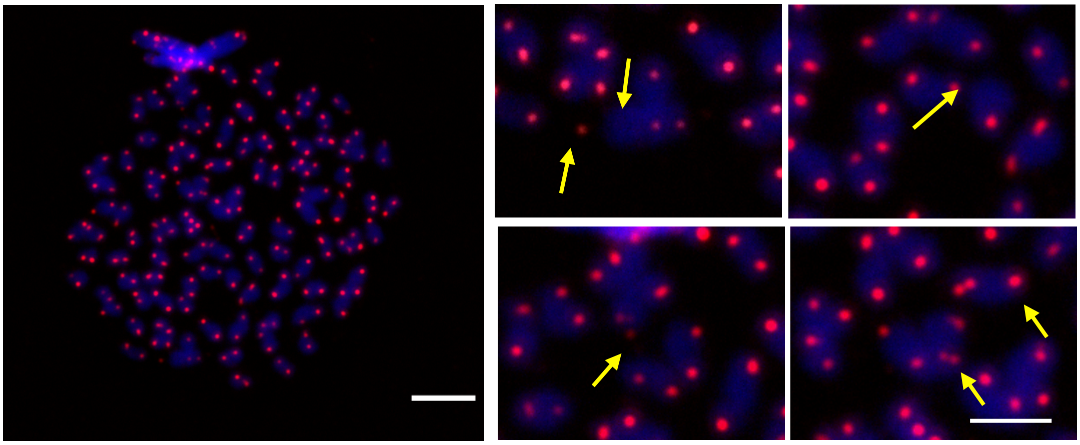

b

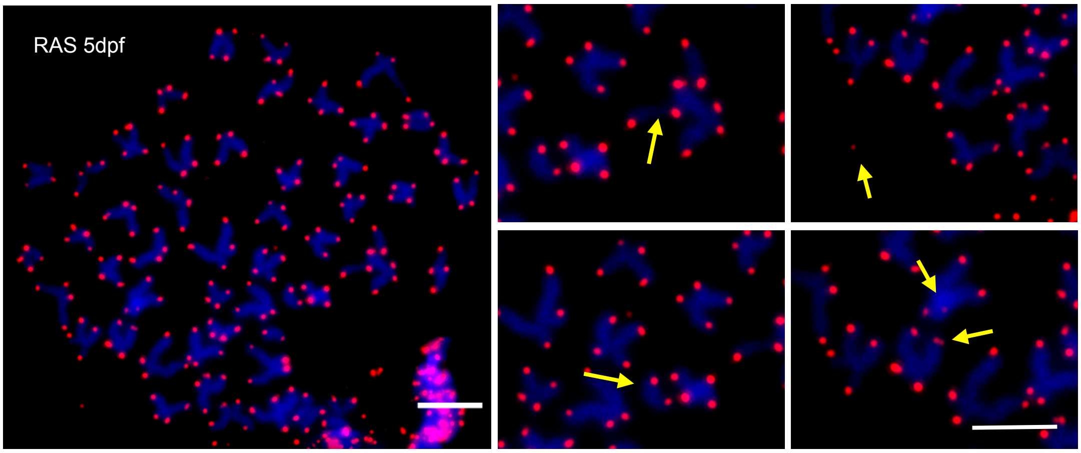


**Figure S3 (related to: A reduction of *tert* expression precedes the development of ALT)**

**a-b)** Representative images of metaphases from zebrafish RAS larvae. Yellow arrows show telomeres abnormalities, similar to those found in adult brain tumors. Scale bars: 1µm.

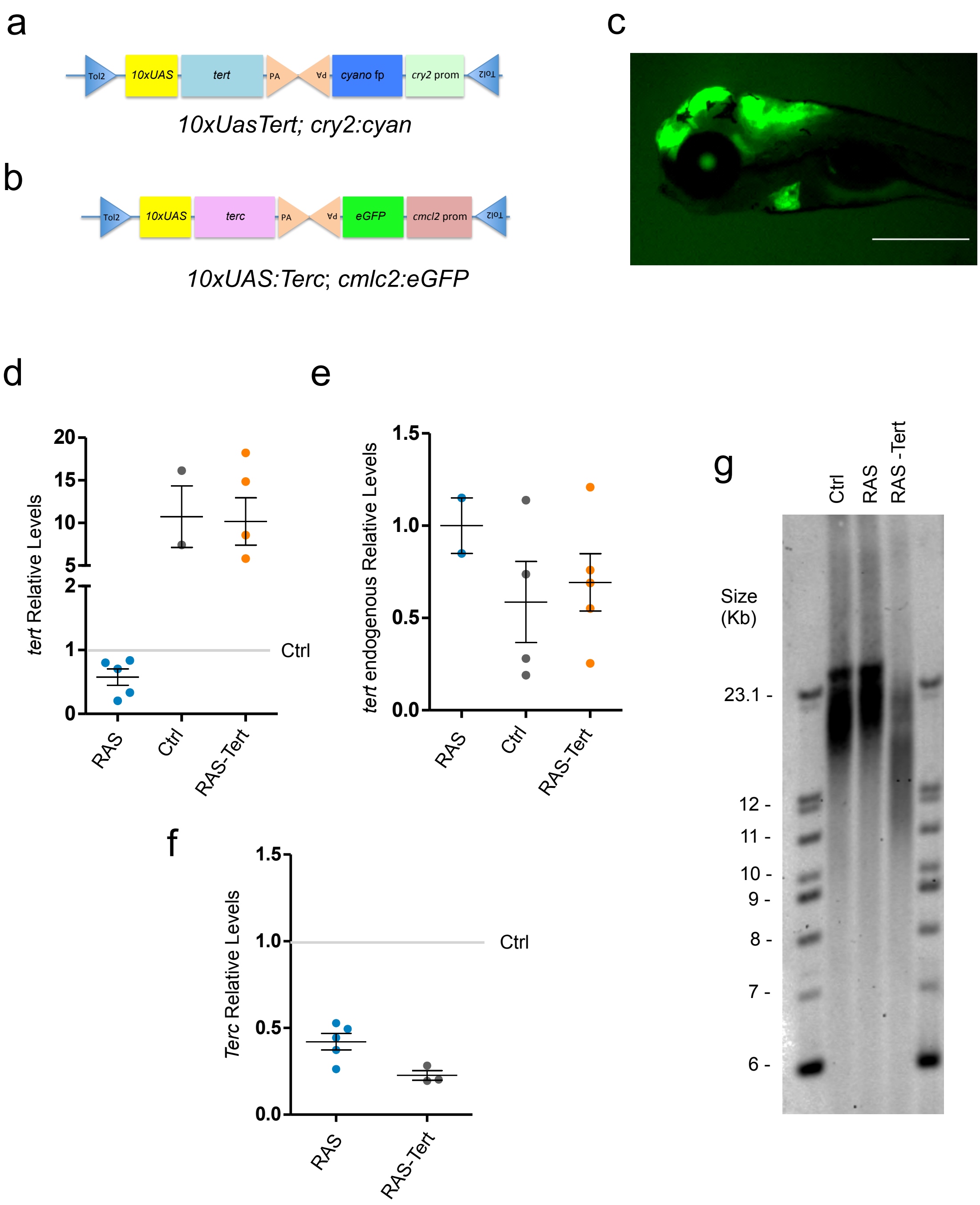


**Figure S4 (related to Figure 6: An overexpression of functional telomerase rescue ALT development)**

**a – b)** Schematic drawing of the plasmids used to generate *tg(10xUAS:Tert)* and *tg(10xUAS:Terc)* transgenic lines. **c)** Image of a 5dpf larva, transgenic for UAS:Tert (green eye), UAS:Terc (green heart) and RAS (green brain). Scale bar:1mm

**d – e – f)** Q-PCR analyses of *tert*, endogenous *tert* and *terc* expression in brain tumors. Values were normalized first to *rps11* and then to total *tert*, endogenous *tert* and *terc* expression in control brains (grey line) (n =2-4).

**g)** Terminal Restriction Fragment analysis of telomeres in Control, RAS and RAS-Tert tumors. A 1.6 kb probe was used to hybridize the membrane.

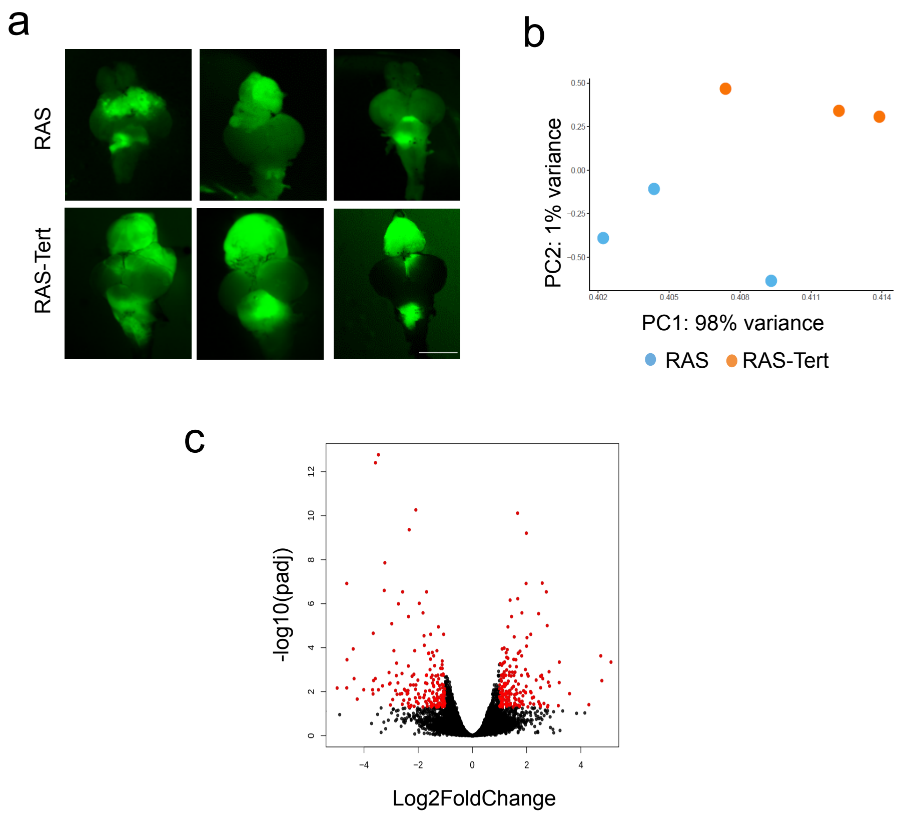


**Figure S5 (related to Figure 7: Molecular differences between ALT+ and telomerase+ brain tumors)**

**a)** Images showing RAS and RAS-Tert brains used for the RNA-Seq analysis. Scale bar: 0.5 mm.
**b)** Principal component analysis (PCA) of the gene expression counts (Trimmed Mean of M-values, TMM) showing the first versus the second principal component (PC). Samples in the two conditions are highlighted in different colors.
**c)** Volcano plot representing -log10 Benjamini-Hochberg (BH) adjusted P-value and log2 Fold Change of genes in the comparison RAS and RAS-TERT. In red differentially expressed genes with BH adjusted P-value <0.05 and log2 Fold Change >1 or <-1 are highlighted.

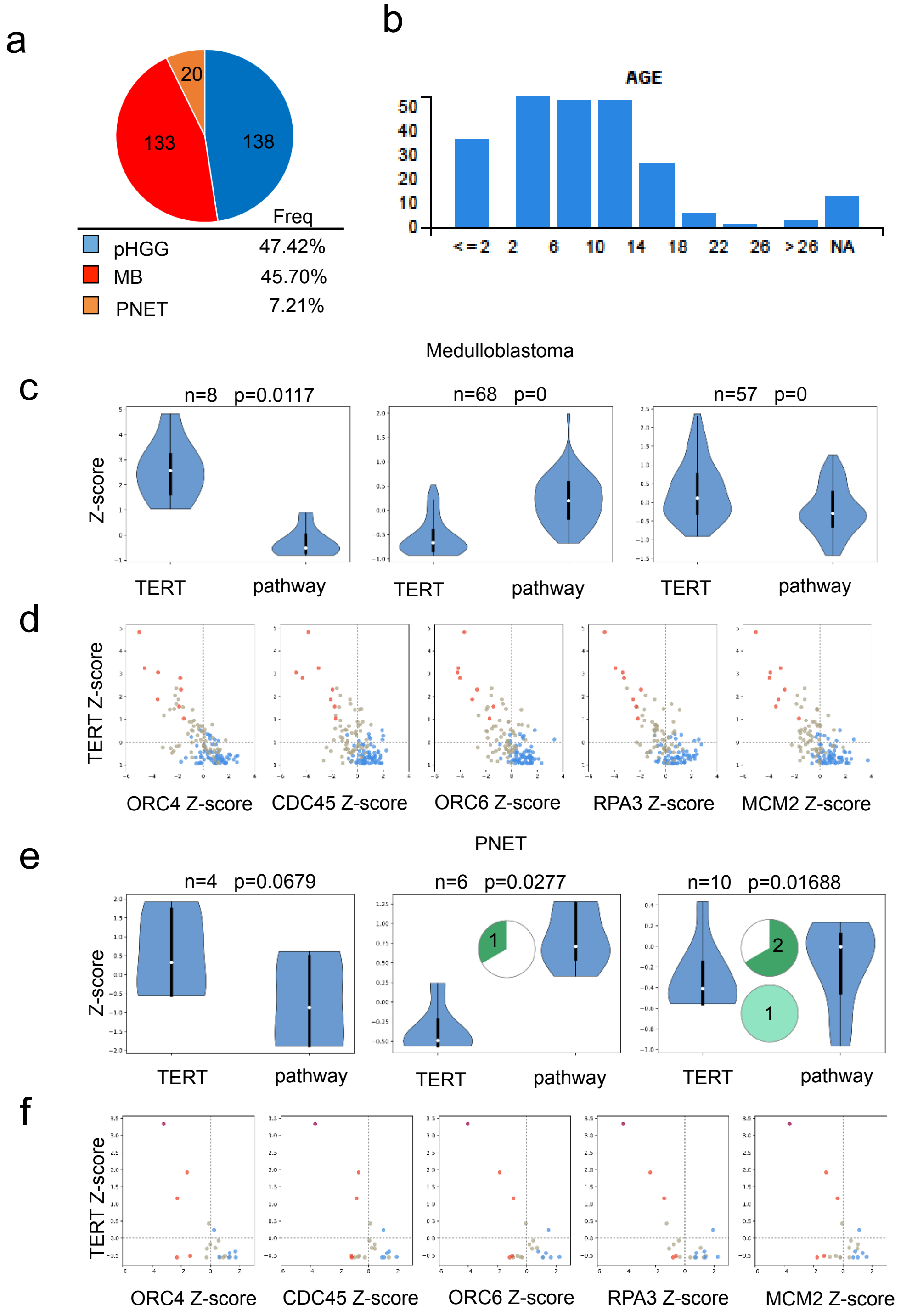


**Figure S6. (related to Figure 7: Analysis of RNA-Seq showed a class of genes altered in zebrafish brain tumors with different TMMs)**

**a)** Pie chart depicting the tumor samples from the pediatric cBio Portal used in the analysis shown in Figure 7. **b)** Diagrams showing the different ages of the patients from the pediatric cBio Portal
**c, e)** Violin plots comparing the expression of TERT and the mean expression of genes in the: Activation of the pre-replicative complex pathway. MB and PNET samples were divided in three groups by K-means clustering. Pie charts show the number of samples with H3F3A (dark green) or ATRX (light green) mutations for each of the three groups. No mutational data in MB were present.
**d, f)** Scatter plots of MB and PNET as indicated, showing the expression of TERT (y-axis, in Z-score) vs the expression of five genes (*ORC4, CDC45, ORC6, RPA3 and MCM2*) in the "Activation of the pre-replicative complex" pathway, that are differentially expressed in zebrafish samples (x-axis, in Z-score). Samples marked in different colors belong to the three different groups as clustered by K-means.

**Legends to Table S1 and S2**

**Table S1: Differentially expressed genes revealed by transcriptome analysis.**

The table reports counts normalization to Trimmed Mean of M-values (TMM) revealed by transcriptome analysis in RAS-Tert versus RAS zebrafish brain tumours. 366 differentially expressed genes (DEG) were idendified using DESeq2, considering adjusted P- value <0.05 and a log2 fold change greater than 1 or smaller than -1. Ensemble ID, gene names and description are reported together with baseMean, log2FoldChange, log2FoldChange (lfcSE), pvalue and p adjusted value (padj). The human orthologous of 296 DE genes are also listed with their ensemble ID (Hs_ensembl_gene_) and human gene name.

**Table S2: Reactome pathways analysis**

Pathway analysis performed with Reactome; enriched pathways are listed according to p-value and p adjusted value. The name of the pathways and the identification code are reported, together with the names and the number of DE genes identified by transcriptome analysis in RAS-tert versus RAS zebrafish brain tumours.

### Table S3

| TelNet functions | DE genes | % TelNet DE genes per listed function |
|---|---|---|
| CHROMATIN ORGANIZATION | SGO1, HIST1H4A, EZH1, KMT2C, DPY30, C17orf49, HMGN2, HMGB2 | 36,36 |
| DNA REPLICATION | CDC45, GAPDH, ORC4, RPA3, RECQL5, LIG1, MCM2, ORC6 | 36,36 |
| TELOMERASE ACTIVITY | DHX36, TFAP2C, DCK | 13,64 |
| TERRA | ORC4, ORC6 | 9,09 |
| DNA RECOMBINATION | RPA3, LIG1 | 9,09 |
| PROTEIN SYNTHESIS | FKBP5, UCHL1 | 9,09 |
| CHROMATIN STRUCTURE | HIST1H4A | 4,55 |
| CHROMATID COHESION | SGO1 | 4,55 |
| ALT ASSOCIATED PML NUCLEAR BODIES | PIAS1 | 4,55 |

**Table S3: Classification of DEG identified in brain tumour models based on TelNet specific genes functions.**

The table reports the most representative telomere related-functions of differentially expressed genes revealed by transcriptome analysis between RAS-tert and RAS tumors, found in the TelNet database (http://www.cancertelsys.org/TelNet/). 51 genes were reported into the relative TelNet functions categories, The percentage identified the the most representative functions impaired.

#### Table S4

| Activation of the pre-replicative complex<br>Homo sapiens R-HSA-68962 | |
|---|---|
| **UniProt** | **Genes** |
| Q9UBD | ORC3 |
| O43913 | ORC5 |
| O43929 | ORC4 |
| Q13416 | ORC2 |
| Q9UJA3 | MCM8 |
| Q9Y5N6 | ORC6 |
| Q13415 | ORC1 |
| Q99741 | CDC6 |
| Q9H211 | CDT1 |
| P25205 | MCM3 |
| P33991 | MCM4 |
| P33992 | MCM5 |
| Q14566 | MCM6 |
| P33993 | MCM7 |
| P49736 | MCM2 |
| O75496 | GMNN |
| P56282 | POLE2 |
| Q07864 | POLE |
| Q9NRF9 | POLE3 |
| Q9NR33 | POLE4 |
| Q9UBU7 | DBF4 |
| O00311 | CDC7 |
| Q7L590 | MCM10 |
| P24941 | CDK2 |
| O75419 | CDC45 |
| Q13156 | RPA4 |
| P15927 | RPA2 |
| P35244 | RPA3 |
| P27694 | RPA1 |
| P49642 | PRIM1 |
| P09884 | POLA1 |
| Q14181 | POLA2 |
| P49643 | PRIM2 |

**Table S4: List of genes in the "Activation of the pre-replicative complex" Reactome pathway**

List of the genes of the: Activation of the pre-replicative complex pathways - Homo sapiens, R-HSA-68962, identified by Reactome analysis using the 366 differentially expressed genes between RAS-Tert versus RAS tumours. This list of genes was used to analyse the activation of the pathway, in paediatric brain tumors data retrieved from pedCbioPortal (https://pedcbioportal.org/login.jsp). UniProt Genes code and gene names are reported.

### Table S5

| Histological code | Code | Tumor Type | Additional features | Age (y/o) | Ccircles dotblot | Ccircles qPCR | TERT | PMLbodies | ATRX |
|---|---|---|---|---|---|---|---|---|---|
| 12913/02 A1 | # 1 | **Medulloblastoma, Classic** | - | 3 | ≈ | + | 0 | 1 | + |
| 15553.3/06 A1 | # 2 | **Medulloblastoma, Classic** | - | 32 | + | ≈ | 0 | 2 | + |
| 8151/00 A3 | # 3 | **Medulloblastoma, Desmoplastic/Nodular** | - | 1 | - | ND | 1 | 0 | + |
| 15189/01 A6 | # 4 | **Medulloblastoma, Desmoplastic/Nodular** | - | 4 | ≈ | + | 0 | 1 | + |
| 12296/07 1G (VI) | # 5 | **Central Nervous System Primitive NeuroEctodermal Tumors (CNS-PNET)** | mostly undifferentiated | 3 | - | + | 0 | 0 | NE |
| 8306.2/01 (PD) | # 6 | **Central Nervous System Primitive NeuroEctodermal Tumors (CNS-PNET)** | mostly undifferentiated | ND | + | - | 0 | 2 | NE |
| 11106/01 (PD) | # 7 | **Central Nervous System Primitive NeuroEctodermal Tumors (CNS-PNET)** | mostly undifferentiated | ND | + | + | 0 | 1 | NE |
| 22994/08 1B (VI) | # 8 | **Central Nervous System Primitive NeuroEctodermal Tumors (CNS-PNET)** | with areas of neuronal differentiation | ND | - | - | 0 | 1 | NE |
| 18345/14 1A | # 9 | **Central Nervous System Primitive NeuroEctodermal Tumors (CNS-PNET)** | with areas of neuronal differentiation | ND | - | + | 0 | 1 | NE |
| 18689.2/16 K1 | # 10 | **Anaplastic Oligodendroglioma** | - | 44 | + | + | 0 | 2 | + |
| 930.1/15 A6 | # 11 | **Anaplastic Astrocitoma** | - | 59 | - | - | 0 | 0 | + |
| 15152/11 | # 12 | **Glioblastoma** | - | 61 | - | - | 1 | 0 | + |
| 24181.1/16 K1 | # 13 | **Glioblastoma** | - | 21 | ≈ | ≈ | 2 | 0 | + |
| 20424.1/16 A1 | # 14 | **Glioblastoma** | - | 50 | - | - | 2 | 0 | + |
| 15998.1/16 A3 | # 15 | **Glioblastoma** | - | 49 | - | - | 1 | 0 | + |
| 39060.2/16 A4 | # 16 | **Glioblastoma** | mostly undifferentiated | 33 | + | + | 3 | 3 | + |
| 27931.1/16 A3 | # 17a | **Glioblastoma with Primitive Neuronal Component (GBM-PNC), IDH1-R132H mutated** | glioblastoma component | 27 | ++ | ++ | 2 | 2 | - |
| A1 | b | | primitive neuronal component | | ≈ | ≈ | 0 | 0 | - |
| 32563.1/15 A6 | # 18a | **Glioblastoma with Primitive Neuronal Component (GBM-PNC)** | glioblastoma component | 76 | + | ND | 2 | 1 | + |
| A5 | b | | primitive neuronal component | | + | ≈ | 0 | 0 | + |
| 36357/15 A2 | # 19 | **Glioblastoma with Primitive Neuronal Component (GBM-PNC)** | glioblastoma component | 68 | ND | ND | 3 | 1 | + |
| A2 | | | primitive neuronal component | | ND | ND | 0 | 0 | + |
| 6622.2/15 A3 | # 20 | **Glioblastoma with Primitive Neuronal Component (GBM-PNC)** | glioblastoma component | 50 | ND | ND | 1 | 1 | + |
| A3 | | | primitive neuronal component | | ND | ND | 0 | 0 | + |

| SCORE | Combined % and intensity |
|---|---|
| very low/negative | 0 |
| low | 1 |
| high | 3 |

**Table S5: Identification of TMMs in a panel of twenty human pediatric brain tumours.**

The table reports features of the 20 brain tumors analysed for C-Circles (by dot blot and Q-PCR), TERT, PML bodies, and ATRX protein expression (by immunohistochemistry). For each tumor, the table reports the histological code, the internal number code (this paper), patients' age and additional features, according to Louis et al., 2016. Colors of highlights represent the TMM classification, and are as follow: Green: ALT; Dark pink: ALT with TERT expression; light pink: ALT with no C-Circle detection; light blue: Telomerase+; not highlight means no TMM classification. Scores are reported in the legend. ND= not detected; NE = not evaluated
